# Linear and categorical coding units in the mouse gustatory cortex drive population dynamics and behavior in taste decision-making

**DOI:** 10.1101/2025.10.06.680705

**Authors:** Liam Lang, Camelia Yuejiao Zheng, Jennifer M Blackwell, Giancarlo La Camera, Alfredo Fontanini

## Abstract

Cortical circuits produce time-varying patterns of population and single neuron activity that play a fundamental role in perceptual and behavioral processes. However, the functional contributions of individual neuron activity to population dynamics and behavior remain unclear. Here we addressed this issue focusing on the mouse gustatory cortex (GC) and using a taste mixture-based decision-making task, high-density electrophysiology, and computational modeling. GC population dynamics represented stimuli linearly during taste sampling and choices categorically before decisions. Single neurons were classified by their linear and categorical activity patterns, revealing subpopulations encoding sensory, perceptual, and decisional variables. To test their functional role, we built a recurrent neural network model of GC. Model perturbations showed linear and categorical neurons were essential for driving normal population dynamics and behavioral performance, whereas many units with other activity patterns could be silenced without consequence. These results have implications that extend beyond GC, and demonstrate the role of linear and categorical coding neurons in cortical dynamics and behavior during perceptual decision-making.

## Introduction

Cortical circuits produce time-varying patterns of population and single neuron activity that play a fundamental role in perceptual and behavioral processes (***Shuler and Bear, 2006; Guo et al., 2014b; Buetfering et al., 2022***). Over the past decade, the gustatory cortex (GC) has emerged as a model for studying such cortical dynamics (***Arieli et al., 2022; Livneh et al., 2020; Mahmood et al., 2025; Vincis et al., 2020; Vincis and Fontanini, 2016***). Population and single neuron activity in GC show characteristic time-varying modulations encoding multiple variables associated with a gustatory experience. Early studies on GC dynamics provide evidence for three temporal epochs following intraoral delivery of tastants in rats. Right after taste delivery, GC neurons encode the somatosensory component of the stimulus hitting the tongue. After a few hundred milliseconds, neurons begin to encode the chemical identity of the tastant and, ultimately, its palatability (***Katz et al., 2001***). A similar sequence of activity has been described also in the GC of mice actively sampling tastants by licking a spout (***Bouaichi and Vincis, 2020***). These dynamics are not limited to the processing of taste; GC can also represent signals related to expectation (***Mazzucato et al., 2019; Livneh and Andermann, 2021***) and decision-making (***Vincis et al., 2020; Lang et al., 2023***).

Recent work relying on delayed response decision-making paradigms shows that populations as well as single neurons in GC can sequentially encode sensory, perceptual, and decisional variables. Head restrained mice were trained in a two-alternative choice (2AC) task (***Guo et al., 2014a; Churchland and Ditterich, 2012***) to sample gustatory stimuli, wait during a delay period, and lick either a left or a right spout based on specific taste-direction associations. In the context of this task, GC activity progresses from taste-coding during the sampling period to representing the licking direction predicted by each taste during the delay period (***Vincis et al., 2020; Lang et al., 2023***). Consistent results were observed in a similar task relying on taste mixtures to guide directional licking decisions. Two-photon calcium imaging has revealed that GC activity first encodes mixture components linearly and then, during the delay period, represents the binary decision to lick left or lick right (***Kogan and Fontanini, 2024***).

These dynamics are not epiphenomenal, as perturbations of neural activity at specific times in the trial interfere with ingestive behaviors as well as taste-guided decision-making. For instance, silencing GC during the palatability epoch delays the onset of aversive reactions to taste (***Mukherjee et al., 2019***). In a 2AC task, silencing GC during the delay period affects task performance (***Vincis et al., 2020***). The importance of these dynamics has therefore spurred extensive investigations of their underlying mechanisms. Spiking network models have unveiled architectural features that are sufficient for producing population dynamics matching those seen in GC during taste-processing, expectation, and decision-making (***Miller and Katz, 2010; Mazzucato et al., 2015, 2016, 2019; Lang et al., 2023***). Furthermore, these studies have provided important information on how perturbing activity during different temporal epochs can impact behavior (***Lang et al., 2023***). Yet, as important as this work has been in advancing our understanding of cortical dynamics, many questions remain unanswered. The relationship among individual neuron activity patterns, population-level dynamics, and behavioral outcomes has yet to be fully elucidated. Most critically, we still do not understand how neurons encoding specific task features contribute to population dynamics and influence overall performance.

Previous computational studies were not designed to investigate the role of specific single neuron firing patterns as both network connectivity and the contribution of functional groups of neurons were established *a priori* and tuned to obtain the desired dynamics and behavior. In this study we overcome the limitations of previous approaches by combining high-density behavioral electrophysiology (***Jun et al., 2017***) with recurrent neural network (RNN) modeling (***Cohen et al., 2020; Valente et al., 2022***). Specifically, we recorded ensembles of GC neurons in mice performing a taste mixture directional task analogous to the task used by ***Kogan and Fontanini (2024***). Population analyses revealed a progression from linear encoding of taste concentrations to categorical prospective encoding of directional licking decisions. Guided by population dynamics, we identified single units that either encoded taste concentration linearly, or task events categorically. A subpopulation of neurons linearly tracked the concentration of the components in the mixture, while others categorically encoded the prevailing taste quality or the imminent licking direction. Linear coding of the stimulus was more predominant early on, following taste sampling, while categorical coding of licking direction was more pronounced later in the trial, before lateral licking. Categorical coding of taste quality was present, albeit in small percentage, throughout the period from sampling to lateral licking. To test the functional significance of single neuron firing patterns, we trained an RNN on both the recorded neural activity and the behavioral performance of the mice for each session. A fraction of the network’s units were trained to match the firing activity of recorded single units, while the rest of the units were free from constraints during training. All the single units recorded in an individual session were fed to the network to ensure that the RNN would not be biased toward linear or categorical patterns or any experimenter-selected feature. After training, the RNNs exhibited linear and categorical patterns in both their constrained and unconstrained units. We hypothesized that these units with specific activity patterns, though relatively small compared to the overall population, were critical for population activity and behavior. We tested this hypothesis by re-running the networks after systematically removing these units that had emerged during training. The simulations revealed that linear coding neurons as well as categorical neurons representing both perceptual and decisional variables were indeed necessary for driving realistic population dynamics and behavioral performance, while a large fraction of the units, exhibiting different patterns of activity, could be silenced without consequence for performance.

Altogether, the results presented here demonstrate the functional significance of specific single neuron firing patterns observed in GC during a taste mixture 2AC task and establish an approach that successfully relates population-level dynamics, individual neuron activity patterns, and behavioral outcomes. Furthermore, since these dynamics are not unique to GC, but have been observed in a variety of cortical circuits (***Goltstein et al., 2021; Niessing and Friedrich, 2010; Reinert et al., 2021***), our work provides insights that generalize far beyond taste.

## Results

### Behavioral task and electrophysiological recordings

Thirteen mice were tested on a binary sucrose/NaCl mixture-based perceptual decision-making task (**Figure 1A**). The task design is based on a two-alternative choice (2AC) paradigm (***Guo et al., 2014a; Churchland and Ditterich, 2012; Kogan and Fontanini, 2024***). Briefly, mice were trained to sample either sucrose (100 mM) or NaCl (100 mM) from a central spout (for ~0.9 s), wait for a delay period (~3.0 s), then lick one of two lateral spouts according to the task policy (e.g., sucrose → lick left; NaCl → lick right). Taste-side pairings were counterbalanced across subjects. Correct lateral licks were rewarded with a drop of water from the lateral spout. The delay period—important for temporally separating the sensory and cognitive processing during this task—was implemented such that the average time between the first lick to the central spout (time point “T”) and the first lick to a lateral spout (time point “D”) was ~3.9 s. Upon reaching criterion for discriminating sucrose from NaCl (i.e., 85% correct performance for three days in a row), mice were tested with the following mixtures (expressed in %Sucrose/%NaCl): 0/100, 25/75, 35/65, 45/55, 55/45, 65/35, 75/25, or 100/0. If the mixture was predominantly NaCl, mice had to lick on the side that was trained to be associated with NaCl; if it was predominantly sucrose, mice had to lick on the sucrose side.

**Figure 1.**
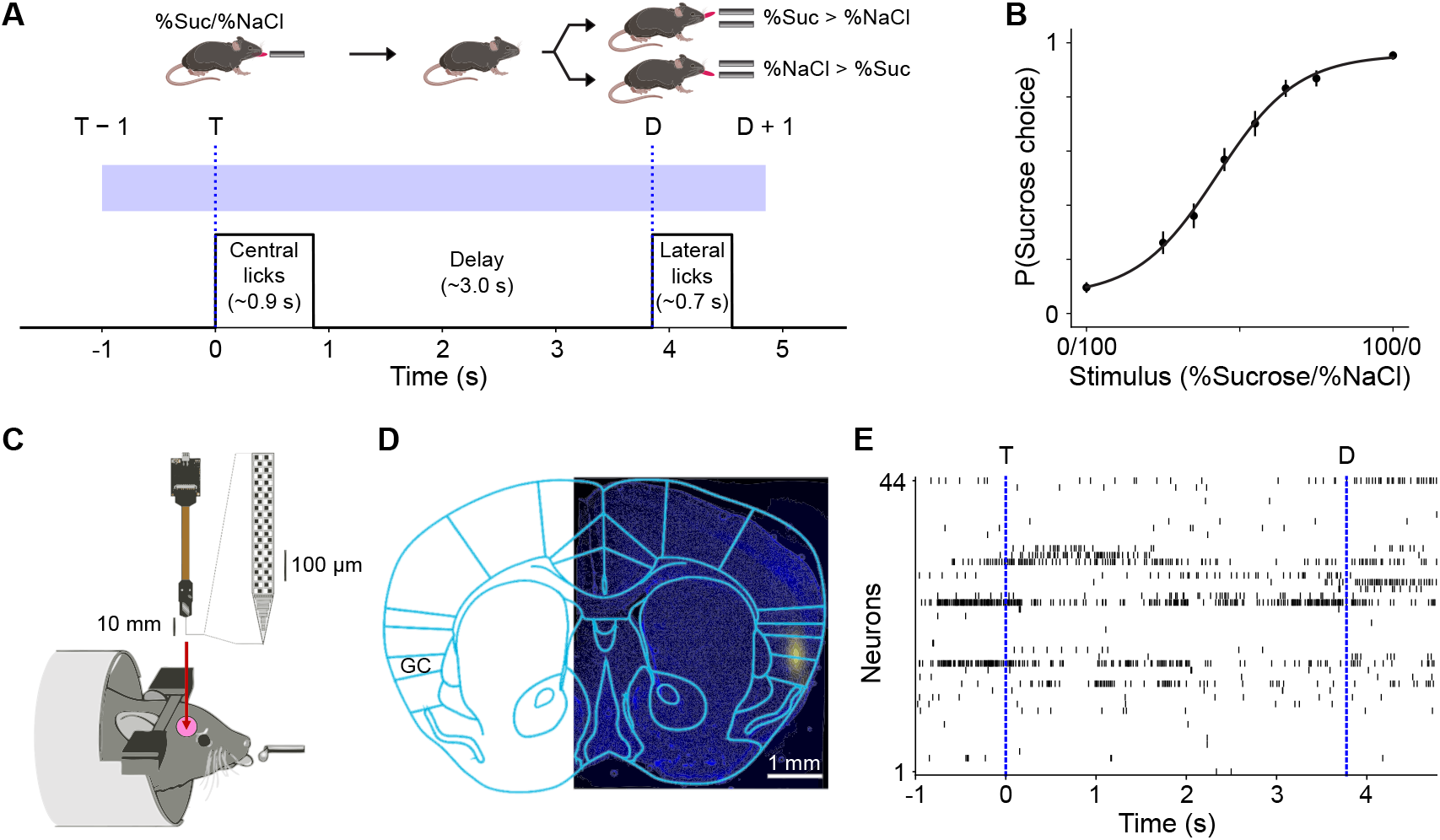
Experimental paradigm. **A**: Schematic of the behavioral task. The blue rectangle indicates the temporal window over which all analyses were conducted, from 1 s before the first central lick, T, to 1 s after the first lateral lick, D. **B**: Psychometric curve averaged across sessions and subjects. Circles and error bars represent mean ± s.e.m. (*N* = 23 sessions across 13 subjects) for the probability of a choice in the sucrose-associated direction for each stimulus value; the continuous curve is a sigmoidal curve fit to the means. **C**: Schematic of acute probe insertion. **D**: Example histological section indicating accurate probe placement in GC (blue: Hoechst; yellow: DiI applied to probe). **E**: Example spike raster plot for neurons simultaneously recorded within a single session from GC of behaving mice.

Mice were tested on this task up to 3 times each (average ~1.8 sessions per animal), resulting in a total of 23 sessions. Mice performed an average of 137 trials per test session (range: 65 to 195) with an average overall accuracy of 77.2 ± 4.3% (range: 69.7% to 86.2%). As expected, performance was near chance level for difficult discriminations (average 56.9 ± 8.9% over sessions for 45/55 and 55/45 trials) and well above it for easy discriminations (average 93.1 ± 3.5% over sessions for 0/100 and 100/0 trials). The psychometric curve for session-averaged performance is plotted in **Figure 1B**. There was a slight asymmetry in the curve (i.e., P(Sucrose choice | Stimulus = 50/50) > 0.5), which could be due to mismatches in perceived intensities of mixture components or to lateral motor biases, but given its small magnitude, we did not investigate it further. Just before each testing session, high-density Neuropixels probes were acutely inserted in the GC so that single unit electrophysiological recordings could be obtained during performance of the test (**Figure 1C-E**). Electrode positioning was reconstructed histologically upon the termination of the experiment. A Python-based GUI for Histological E-data Registration in Brain Space was used to register the slice images onto the Allen CCF mouse atlas (***Fuglstad et al., 2023***). **Figure S1** shows the reconstructed positioning of the probes. All channels were sorted, but only those mapped within GC were used for analysis. Simultaneously recorded GC ensembles had an average size of ~27 neurons (range: 7 to 68), for a total of 626 neurons across all sessions.

### Population dynamics during mixture-based decision-making

To assess the involvement of GC in representing different events within a taste mixture 2AC task, we analyzed task-related, population firing activity. We used a “warped” time scale to allow for consistent analysis of dynamics with respect to multiple behavioral events of interest that have trial-to-trial variability. Each neuron’s peristimulus time histograms (PSTHs) were constructed by aligning spike trains to the first central lick T, warping each trial’s duration between first central and lateral lick to the overall average (~3.9 s), calculating firing rates in ~50 ms bins, averaging over correct trials of each mixture separately, and smoothing with Gaussian kernels. Visual inspection of the population PSTH averaged across all recorded neurons and for all stimuli (**Figure 2A**) revealed increases in firing rates aligned with the central and lateral licks. Consistent with this, we found 57.5% (370/626) of neurons responded to taste and 43.9% (275/626) displayed preparatory activity before lateral licks (see **Methods: Responsivity and selectivity analyses**; **Table S1** provides these counts across sessions). To identify when populations of GC neurons discriminate between trial types (correct predominantly-sucrose vs correct predominantly-NaCl trials), we computed auROCs and measured differences in firing rate distributions between the two types of trials (**Figure 2B**). Neurons could maximally discriminate between correct sucrose and NaCl trial types at all time points (white dots), with noticeable concentrations of differential activity around time points T and D (white traces overlaid on the heatmaps are neuron-averaged auROCs). When separating neurons based on whether their peak differential activity was in favor of predominantly-sucrose or predominantly-NaCl mixtures, we found no significant difference between the number of neurons that “preferred” one to the other (306 vs 320; 2-tailed binomial test, *p* = 0.548). Though there were slight qualitative differences between mean auROC traces, they had similar peaks during sampling (0.062 for NaCl-preferring, 0.052 for sucrose-preferring between T and D) and post-decision (0.077 for NaCl-preferring, 0.075 for sucrose-preferring after D). Moreover, the distribution of peak au-ROCs was not different between NaCl- and sucrose-preferring neurons (rank-sum test, *p* = 0.789).

**Figure 2.**
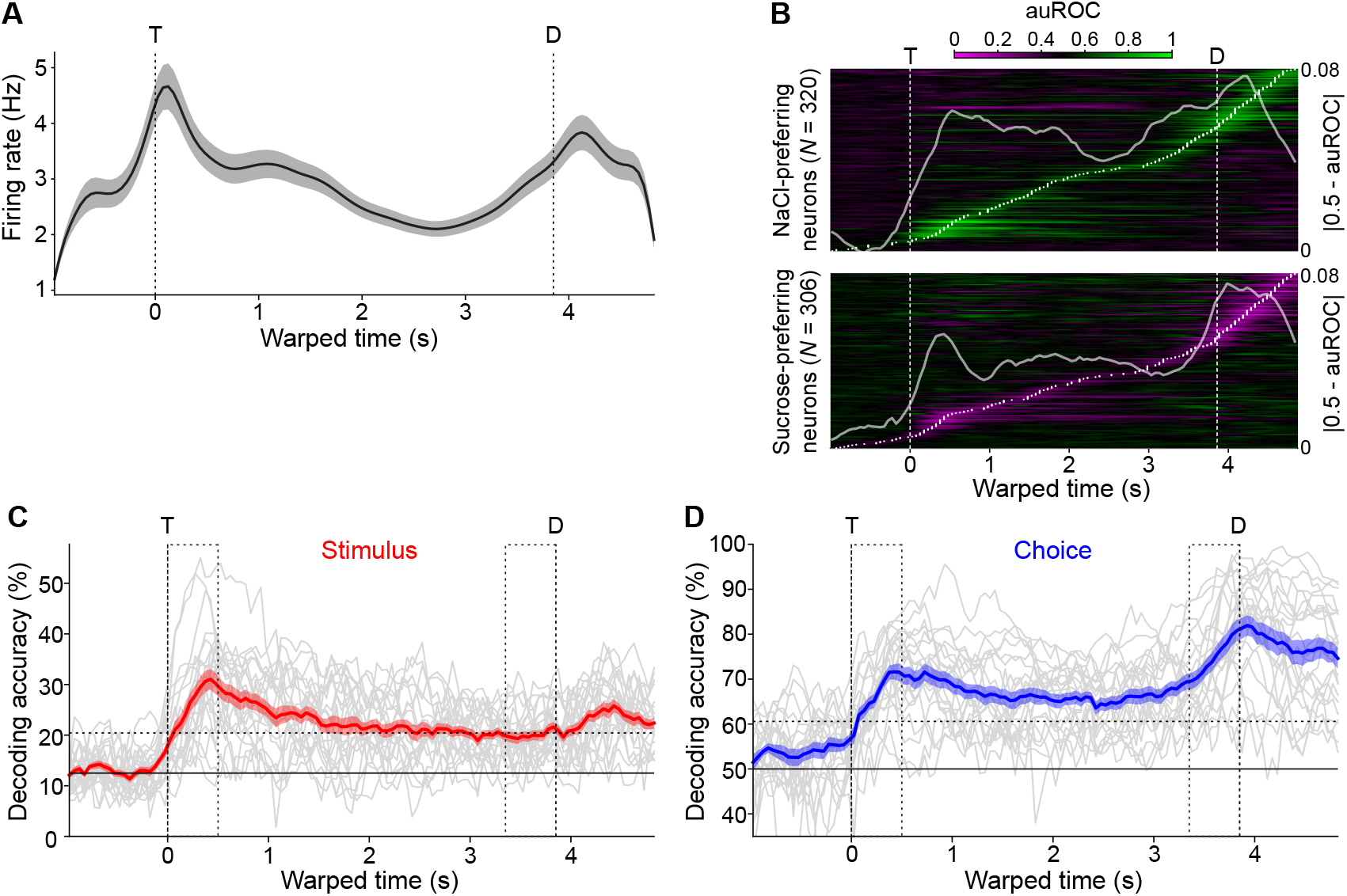
Population activity and information encoding during taste mixture-based decision-making. **A**: Population PSTH (*N* = 626). Vertical dashed lines indicate the first central (T) and lateral licks (D), respectively. The trace represents mean firing rate; shading represents s.e.m. **B**: Population heatmaps for single unit differential activity between correct predominantly-sucrose and correct predominantly-NaCl trials. White dots indicate each unit’s time of peak differential activity. Traces are ordered by peak time and separated by whether peak differential activity is in favor of NaCl (“NaCl-preferring”) or sucrose (“Sucrose-preferring”). White trace is the mean auROC across neurons. **C-D**: Decoding of task-relevant variables. For each session (grey trace), accuracy is plotted over time with colored shaded traces representing the mean ± s.e.m. over sessions. Trial labels to be decoded were mixtures (**C**) and choice (**D**). Horizontal solid line represents theoretical chance level. Horizontal dashed line represents theoretical significant decoding threshold (*α* = 0.01). Dashed rectangles mark the “sampling” and “delay” analysis windows.

To assess how such task-related firing encoded task variables (e.g., stimuli and decisions), we trained classifiers to decode stimulus and choice variables from activity vectors for each ensemble of simultaneously recorded neurons (23 ensembles with a median size of 27 neurons). For each recording session, all trials were labeled according to the delivered stimulus (8 possible labels depending on the mixture) and whether the animal chose the left or right lateral spout. At each time point, a decoder then predicted a label for each trial as the label whose trial-averaged activity vector was nearest (in terms of Euclidean distance) to the activity vector of the trial in question. Session-averaged decoding for both stimulus and choice was significantly above chance throughout the period between taste delivery and reward; however, the dynamics differed significantly (2-way within-subjects ANOVA with factors time [sampling/delay] and decoded variable [stimulus/choice]; interaction *p* < 0.001). Stimulus decoding peaked at 31.0% ~425 ms after stimulus delivery (**Figure 2C**), while choice decoding also rose with stimulus delivery but ramped before the lateral licks and peaked at 81.9% ~75 ms after the decision time (**Figure 2D**).

To further assess response dynamics, population activity trajectories across different trial types were analyzed with both unsupervised and supervised dimensionality reduction techniques. For the unsupervised method, we calculated pairwise Euclidean distances in 626-dimensional neural space between all stimulus types and time points (**Figure 3A**). We then used a t-distributed stochastic neighbor embedding (t-SNE) to non-linearly map the neural data into a 2-dimensional space while attempting to preserve the true distance structure (**Figure 3B**). Notably, activity trajectories diverged at the time of taste delivery according to stimulus type, with more distant stimuli (e.g., 0/100 vs 100/0) corresponding to more distant neural activities and more similar stimuli (e.g., 45/55 vs 55/45) corresponding to closer neural activities. The trajectories then binarized according to stimulus choice, with a large distance between trials predicting different licking directions regardless of exact stimuli. To separate the components associated with the stimulus from those associated with the choice, we employed a demixed principal component analysis (dPCA; ***Kobak et al., 2016***) and found the dimensions of maximal data variance with respect to stimulus- and choice-specific variance. The projection of the population activity onto the principal stimulus component (**Figure 3C**) shows a monotonic, linear separation of activity according to mixture, irrespective of the ultimate choice (note how each average error trial trajectory, the dotted lines, overlaps with its average correct trial trajectory, the solid lines). In contrast, the projection of population activity onto the principal choice component (**Figure 3D**) shows a binary separation of activity according to upcoming choice irrespective of mixture (note how average error trial trajectories for predominantly-sucrose mixtures overlap with average correct trial trajectories for predominantly-NaCl mixtures, and vice versa). The time courses of the projected activities suggest that a binarization of activity according to selected choice emerges just before lateral licking as the graded stimulus-based activity slowly collapses.

**Figure 3.**
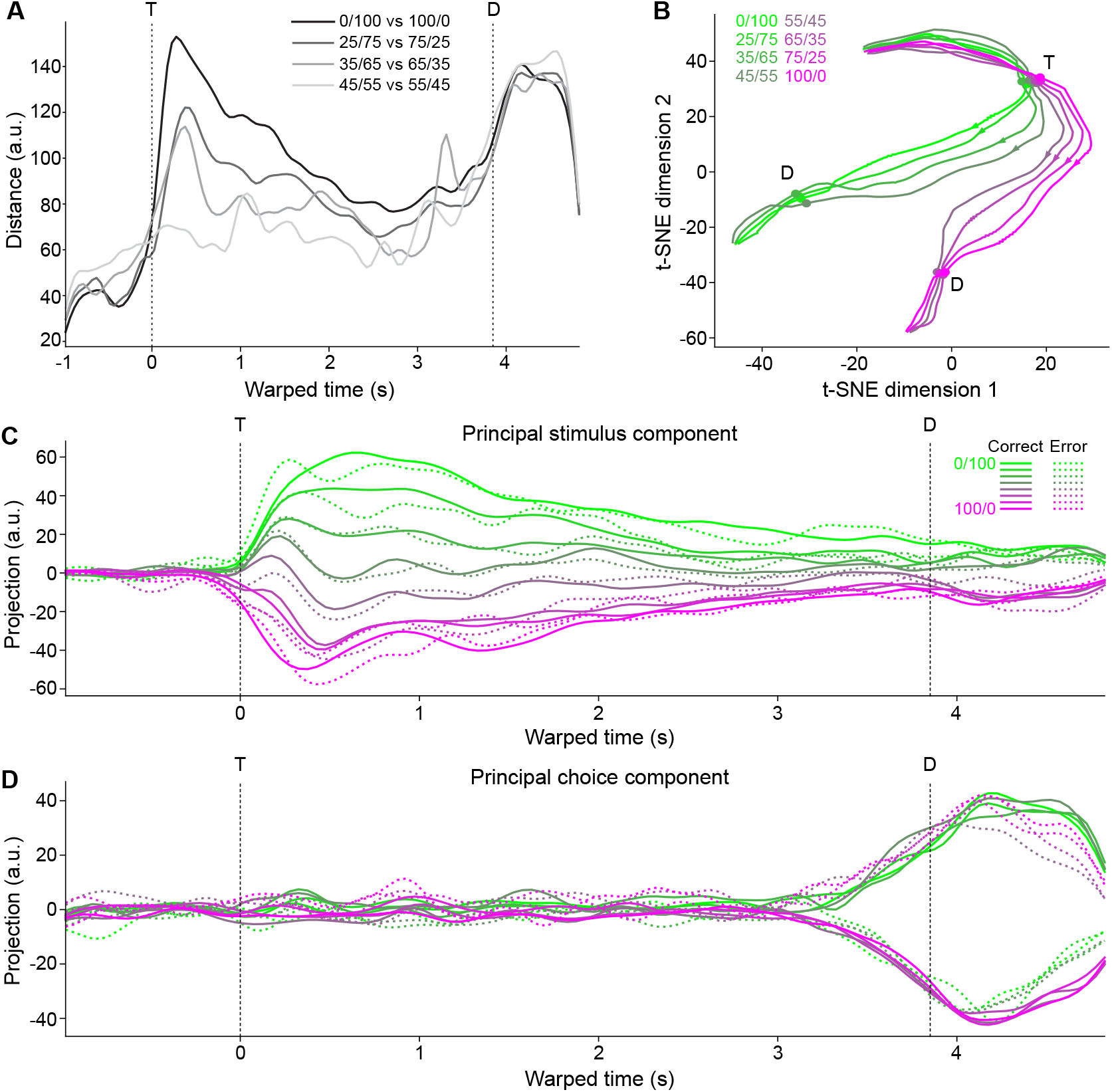
Low-dimensional population activity trajectories. **A**: Euclidean distances between pairs of trial-averaged pseudo-population activity trajectories. **B**: t-SNE of trial-averaged pseudo-population trajectories for all stimuli (%Sucrose/%NaCl) based on pairwise Euclidean distances between activities. **C**: One-dimensional linear projections of trial-averaged pseudo-population trajectories onto the demixed principal component explaining maximum stimulus-specific variance. Solid lines are correct trial averages; dotted lines are incorrect trial averages. **D**: Same as **C**, but for the demixed principal component explaining maximum choice-specific variance. T: time of first central lick; D: time of first lateral lick.

In summary, population analyses show a linear representation of taste mixture information toward the beginning of the trial and a categorical representation of decisions toward the end.

### Single unit responses during mixture-based decision-making

Upon observing that population activity can represent task variables either in a linear or binary fashion, we investigated whether such representations might also be found at the level of single units. A response profile analysis was performed. To extract a response profile (essentially a tuning curve) a neuron’s firing rate in a specified time window was averaged over correct trials of a particular stimulus and plotted as a function of the mixture. Two temporally separated windows of interest, [T, T + 500 ms] and [D – 500 ms, D], were chosen and, for each, the single unit response profiles for all neurons were constructed. Profiles were assigned a label, “linear” or “step”, after a least-squares regression statistically determined the shape that fit the profile best (if neither fit was significant, it was assigned the “other” label; see **Methods: Response profiles**). The shape templates were chosen based on previous published work (***Kogan and Fontanini, 2024; Maier and Katz, 2013***), with linear fits representing response profiles that track the concentration of one of the components in the mixture and step fits representing response profiles that change abruptly at the 50/50 mixture. **Figure 4A** shows examples of linear (left) and step (middle, right) single unit response profiles (bottom), as well as these neurons’ corresponding PSTHs (top).

**Figure 4.**
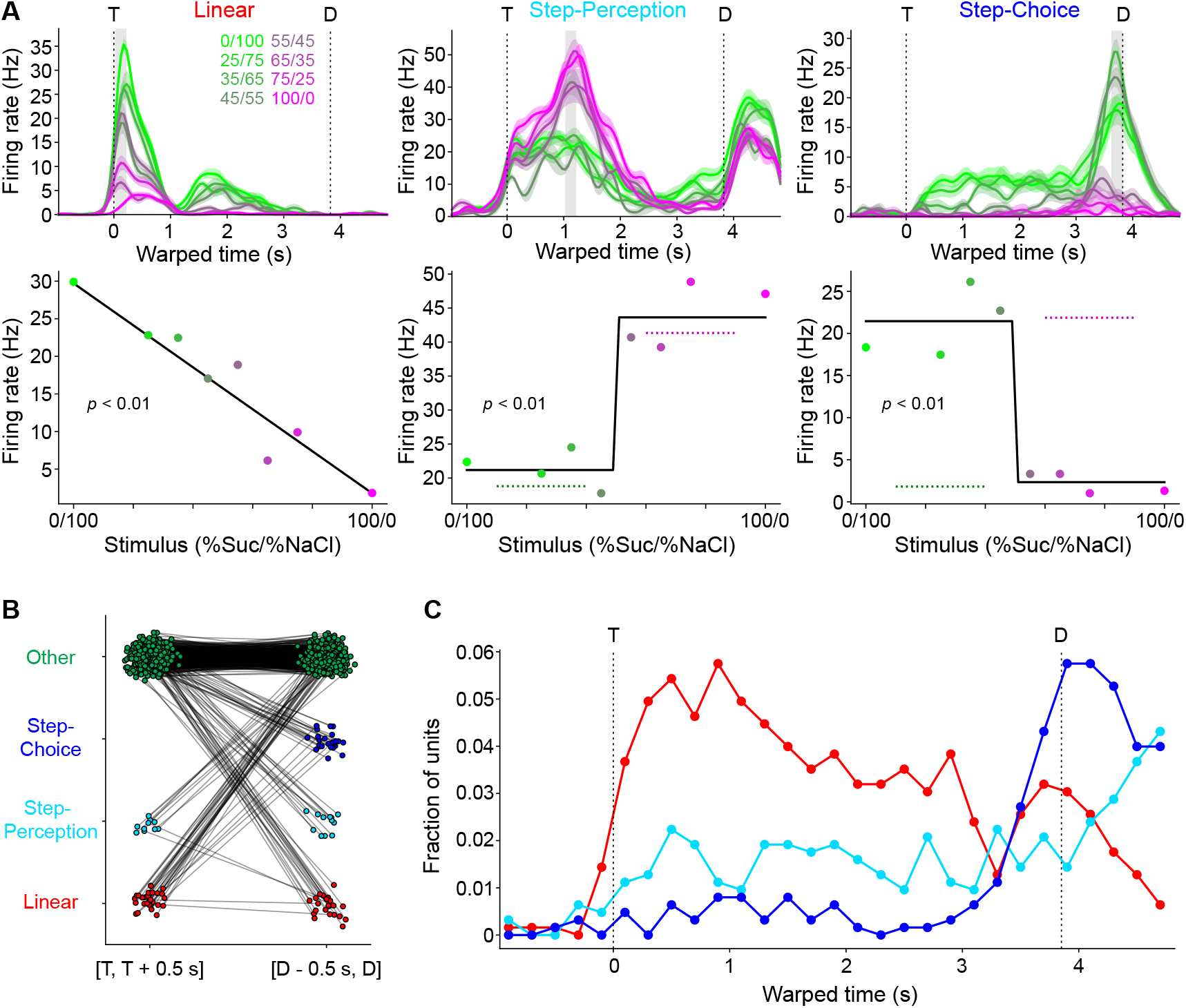
Classification of single unit coding types. **A**: Representative single unit PSTH (top) and response profiles (bottom) exemplifying the different coding types within a time window (grey bar, top): linear (left), step-perception (middle), and step-choice (right). Step-perception (middle) and step-choice (right) types were disentangled by comparing correct trials to error trials (dashed lines in bottom plots). Color scale corresponds to different mixture stimuli (%Sucrose/%NaCl). **B**: Visualization of each neuron’s coding type label (vertical axis) between two time windows (horizontal axis). Each neuron is a point in both windows, with lines connecting the same neurons. T: time of first central lick; D: time of first lateral lick. **C**: Distribution of coding types across all neurons (pooled over all sessions) over time. For each time point (a window ~200 ms wide), the coding type classification analysis depicted in **A** was applied to each neuron.

Step coding neurons could represent either a perceptual category (i.e., sweet vs salty) independently of concentration and licking direction, or the imminent licking direction independently of the stimulus. To disambiguate these options we analyzed error trials—that is, trials in which a mouse received a mixture and licked toward the wrong direction. If a step coding neuron had an average firing rate that was consistently elevated for trials where the same choice was taken, regardless of stimulus, the neuron was defined as a “step-choice” neuron (**Figure 4A**, right). On the contrary, if a step coding neuron had an average firing rate that was elevated for trials where the same stimulus was presented, regardless of the choice, the step coding neuron was considered a “step-perception” neuron (**Figure 4A**, middle).

Comparing the first 500 ms of the sampling period to the last 500 ms of the delay period, there was a similar proportion of neurons whose response profile could be significantly fit by either linear or step functions (6.5%, 41/626 for the first 500 ms vs 9.4%, 59/626 for the last 500 ms, Chi-squared *p* = 0.061) (**Figure 4B**). It is worth noting that neurons with no significant fits could still show significant responses to task events. For instance, in the first 500 ms 57.1% (334/585) were taste responsive and 10.1% (59/585) were taste selective (i.e., they responded differently to the mixtures), while in the last 500 ms 42.0% (238/567) showed preparatory responses in anticipation of lateral licks (63 of which were selective for a specific direction).

Within the sub-groups of neurons with significant fits there was a change in the distribution of response profile types over time: there was a significant increase in the proportion of step coding neurons from the beginning to the end of the trial (24.4%, 10/41 vs 62.7%, 37/59, Chi-squared *p* < 0.001); equivalently, there was a significant decrease in the proportion of linear coding neurons (75.6%, 31/41 vs 37.3%, 22/59, Chi-squared *p* < 0.001). The increase in step coding neurons from the beginning to the end of the trial was driven not by changes in the amount of step-perception neurons, which remained stable (24.4%, 10/41 vs 37.3%, 12/59, Chi-squared *p* = 0.631), but by an increase in the amount of step-choice neurons (0/41 vs 42.4%, 25/59, Chi-squared *p* < 0.001) (**Figure 4B**). Furthermore, the change in distribution over time seemed to be driven mostly by separate populations of linear and step coding neurons, rather than a single population of coding neurons that switched its coding type: 77.4% of the linear coding neurons at the beginning of the trial had no significant response fit at the end of the trial; similarly, 86.5% of step coding neurons at the end of the trial had no significant fit at the beginning of the trial. The remaining neurons multiplexed across time.

To investigate the dynamics of single neuron responses, the time course of the distribution of fits was computed by running the response profile analysis with a moving window of ~200 ms. When quantified in this way, we found that 24.8% (155/626) of neurons were linear coding in at least one bin, 26.7% (167/626) were step-perception, and 18.2% (114/626) step-choice (see **Table S2** for session-by-session data). That said, the peak percent of coding units in any specific bin was much lower (maximum 10.7% total). Visual inspection of **Figure 4C** indicates the same trend of switching from mostly linear coding to mostly step-choice coding over time (**Figure 4C**) seen at the population level (**Figure 3**). In addition, the analysis reveals a small but consistently present proportion of step-perception neurons.

Altogether, single neuron analyses confirm the population results on coding dynamics and extend those findings to show that GC single units can encode in a binary fashion both the choice of licking direction (i.e., left vs right) as well as the perceptual category (i.e., sweet vs salty). These results also raise questions about the functional significance of the relatively low percentage of linear and step coding neurons and their contribution to population dynamics and task performance.

### Recurrent neural networks capture experimental neural and behavioral results

To investigate the functional role of single neuron response types described above, we relied on a computational approach and, for each recording session, built a recurrent neural network (RNN) constrained by the single neuron data and capable of reproducing behavioral performance. For each experimental session, we modeled the simultaneously recorded neurons as a fraction of a larger system of units that received external stimulus input, noise input, and recurrent input from other units. The model was partitioned into units that were trained to reproduce the neural activity directly observed during the experiment, termed “constrained,” and those that were not, termed “unconstrained.” The ratio of total units to constrained units was fixed at 5.88 (thus, constrained units were ~17% of each network). An additional external unit allowed for the model to produce “choice activity” by weighting the firing rates of all internal units. Discrete choices were obtained by thresholding the average choice activity over a decision window (from D – 100 ms to D, for D = 3.9 s, the average decision time in the experimental dataset). The network’s training incorporated two processes: reproducing the experimentally observed patterns of neural activity within the constrained population, while simultaneously selecting the appropriate behavioral response to each stimulus. That is, the output of each constrained unit was trained to match the PSTHs of a corresponding neuron actually recorded from the mouse GC during behavior. This approach, like the one described in ***Cohen et al***. (***2020***), enhanced the biological realism of the RNN trained to perform the task since its internal activity was explicitly instructed to resemble true neural activity. **Figure 5A** illustrates the key components of our RNN model.

**Figure 5.**
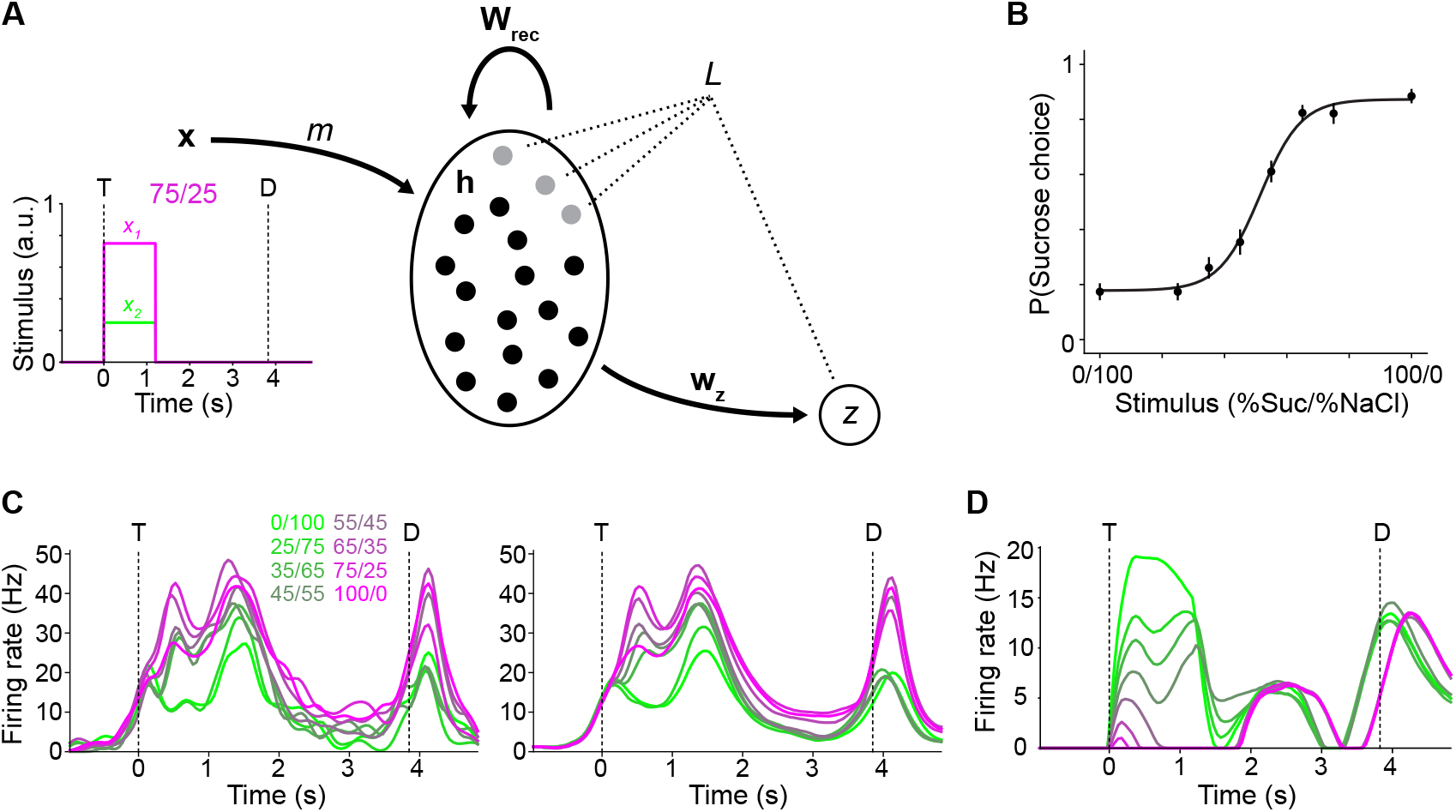
Recurrent neural network design and behavior. **A**: Model architecture. *N* neurons are modeled as dynamic units with internal activity **h** that is influenced by the external stimulus input (*m*(**x**); the time course of an example **x** for mixture stimulus 75/25 is shown), recurrent input (via **W**_rec_), and noise input (not shown). A decision unit *z* measures the network’s choice by taking a weighted sum of activities via **w**_*z*_. The loss function *L* is minimized during training based on choice (*z*) and the activity of the constrained units (grey dots). T: time of stimulus onset; D: decision time. **B**: Psychometric curve fit to across-model means for the probability of the sucrose choice as a function of the stimulus. Circles and error bars represent mean and s.e.m. **C**: Example of experimentally observed PSTH (left) and the corresponding firing rate activity for the unit in the network trained to match it (right). **D**: Example firing rate activity for a unit in the network not explicitly trained to match any experimentally observed PSTH. Color scale corresponds to different mixture stimuli (%Sucrose/%NaCl).

The models successfully learned to perform the decision-making task and produced psychometric curves qualitatively similar to real animals’ when presented with noisy mixture stimuli (**Figure 5B**). Although the psychometric functions were significantly different between model and experiment (extra-sum-of-squares F-test, *p* < 0.001), with a greater slope for the model’s (0.15 vs 0.08), this was unsurprising given that we tuned the level of input noise only to match overall accuracy, which was 77.2%, comparable to the 77.2% we saw from mice (t-test, *p* = 0.964). At the same time, the models successfully reproduced the experimentally observed neural activity. **Figure 5C** shows an example experimental PSTH (left) and the activity of the corresponding unit in the model trained to reproduce it when presented with noiseless mixture stimuli (right). This unit has a root-mean-squared-error between model output and target PSTH of 1.86 Hz; the median value across all constrained units was 1.26 Hz. Additional examples of constrained units are provided in **Figure S2**. Unconstrained units learned to produce a variety of responses to stimuli, some of which resembled patterns seen in the experimental dataset. An example unconstrained unit is shown in **Figure 5D**, and breakdowns of all responses are shown in **Figures S3-4**. In total, 49.8% (1832/3681) of units were taste responsive and 40.8% (1502/3681) of units showed preparatory activity leading up to decisions (see **Table S3** for across-session counts), comparable to the experimental findings.

Additional support for the overall agreement between experimental and model dynamics and coding schemes came from population analyses. 160 trials (20 per mixture stimulus) of data from each model were simulated, using the same levels of noise added to stimuli as were used to produce realistic psychometric curves. We then pooled all the units (constrained and unconstrained) across models and applied the same dPCA procedure we used on experimental data to find the principal stimulus- and choice-coding components. Population activity projected on the principal stimulus component (**Figure 6A**) showed a graded separation of trajectories according to stimulus and regardless of choice, while the projection on the principal choice component (**Figure 6B**) showed a binary separation of trajectories according to choice and regardless of stimulus. The time courses suggested a transition from stimulus-representing activity to choice-representing activity over the period between central and lateral licks. These patterns of activity are qualitatively consistent with those observed in the experimental data and demonstrate that the RNNs produce biologically realistic population dynamics.

**Figure 6.**
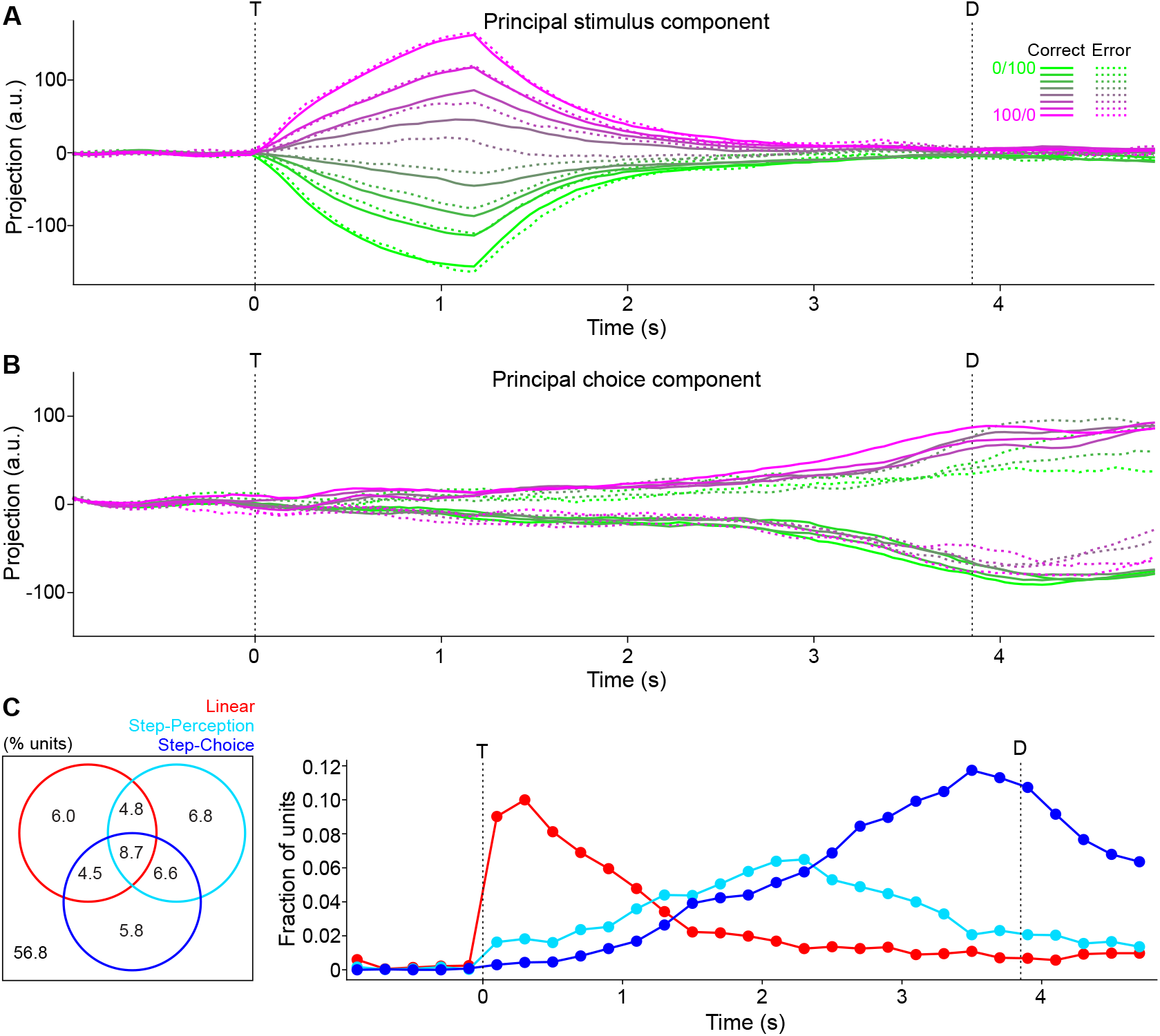
Modeled population activity and single unit coding properties. **A**: Trial-averaged pseudo-population activity trajectories projected onto demixed principal component of maximal stimulus-specific variance. Solid lines are correct trial averages; dotted lines are incorrect trial averages. Color scale corresponds to different mixture stimuli (%Sucrose/%NaCl). T: time of stimulus onset; D: decision time. **B**: Same as **A** but for the demixed principal component of maximal choice-specific variance. **C**: Left: Venn diagram showing percentages of neurons with all possible combinations of coding types over time. Right: Distribution of coding types across all units (pooled over all models) over time.

To further validate the realism of the network and identify the salient features of neural activity on a single unit level, the same response profile classification analysis performed on experimental neurons was applied to model units. Using the results of the simulations described above, response profiles were calculated for all units (for correct and error trials, separately) in 200 ms bins, and the same procedure was used to assign a coding type label—linear, step-perception, step-choice, or “other”—to each response profile in each bin. The Venn diagram in **Figure 6C** (left) displays the breakdown in percentages of all units that exhibited each possible combination of coding types in at least one bin during the task period. In total, 24.2% (885/3681) of units were classified as linear coding, 26.8% (987/3681) of units were step-perception coding, and 25.5% (943/3681) of units were step-choice coding at some time point (see **Table S4** for session-by-session data). As in the case of the experimental results, the model’s neurons that did not show significant fits (56.8% of units) could still be taste responsive (29.4%, 615/2092), taste selective (9.4%, 197/2092), and show preparatory responses (23.4%, 490/2092) (**Figures S3-4**).

Analyses on the time course of the distribution of coding types—linear, step-perception, and step-choice—across all units (pooled over all 23 models) showed qualitative agreement with the experimental findings (**Figure 6C**, right). As in the experimental data, the peak percent of coding units in any specific bin was relatively low (maximum 14.9% total). The prevalence of linear coding units peaked soon after mixture sampling, while the step-choice coding units peaked soon before lateral licking. As in the experimental data, the model produced a lower percentage of step-perception coding units whose prevalence tiled the entire trial. The results of **Figure 6** held even when the analyses were restricted to either the constrained (**Figure S5**) or unconstrained (**Figure S6**) units, with the constrained results qualitatively appearing even more similar to the experimental ones, as expected, and the unconstrained results appearing nearly indistinguishable from the full model results.

Altogether, the results show that the RNN models trained to reproduce behavioral performance and, in a fraction of the units, the observed neural activity, generate population and single unit coding patterns analogous to those observed in GC of behaving mice.

### Model perturbations reveal behavioral significance of coding unit types

Generating a series of RNNs that aligned with experimentally observed neural activity and behavioral performance allowed for an exploration of the functional role of the different types of coding units (linear, step-perception, and step-choice). A series of virtual “ablation” experiments were conducted by re-running simulations while clamping the firing rates of specific sub-populations of units to 0. “Ablation” experiments were conducted for units that exhibited linear coding at any time point in the original simulations, those that were classified as step-perception coding at any time point, those that were labeled as step-choice at any time, and those that never followed any of these coding patterns (i.e., the “other” units).

All three coding types contributed to model dynamical activity along the stimulus- and choice-coding dimensions, as projecting the post-ablation activity onto the originally identified axes resulted in a noticeable blunting (**Figure 7A**). Quantitatively, mean absolute projection values were significantly reduced (post-hoc Bonferroni-corrected paired t-tests; stimulus projections: control vs linear, *p* < 0.001; control vs perception, *p* < 0.001; control vs choice, *p* < 0.001; choice projections: control vs linear, *p* < 0.001; control vs perception, *p* < 0.001; control vs choice, *p* < 0.001). Similarly, new stimulus- and choice-coding dimensions identified after ablating did not align with the old ones (**Figure 7B**) (absolute cosine similarities between vectors; stimulus components: control vs linear, 0.208; control vs perception, 0.244; control vs choice, 0.557; choice components: control vs linear, 0.016; control vs perception, 0.169; control vs choice, 0.021). The effects of the “ablations” could simply be the result of the removal of roughly a quarter of the units in a highly recurrent network leading to a large non-specific disruption of dynamics. However, this is not the case, as the “ablation” of the “other” units (i.e., those that do not show any significant fit), which constitute a much larger fraction of the network (56.8%, 2092/3681), left dynamics largely intact. Activity projections onto the original stimulus- and choice-coding dimensions after ablating “other” units (**Figure 7A**) were similar to the control condition without ablation (**Figure 6A-B**) (stimulus projection: *p* = 0.105; choice projection: *p* > 0.999) and newly identified coding dimensions (after the ablations) overlapped highly with the originals (i.e., before the ablations; **Figure 7B**) (stimulus component: 0.947; choice component: 0.959). This is relevant as neurons in this group show firing modulations to task events (see above).

**Figure 7.**
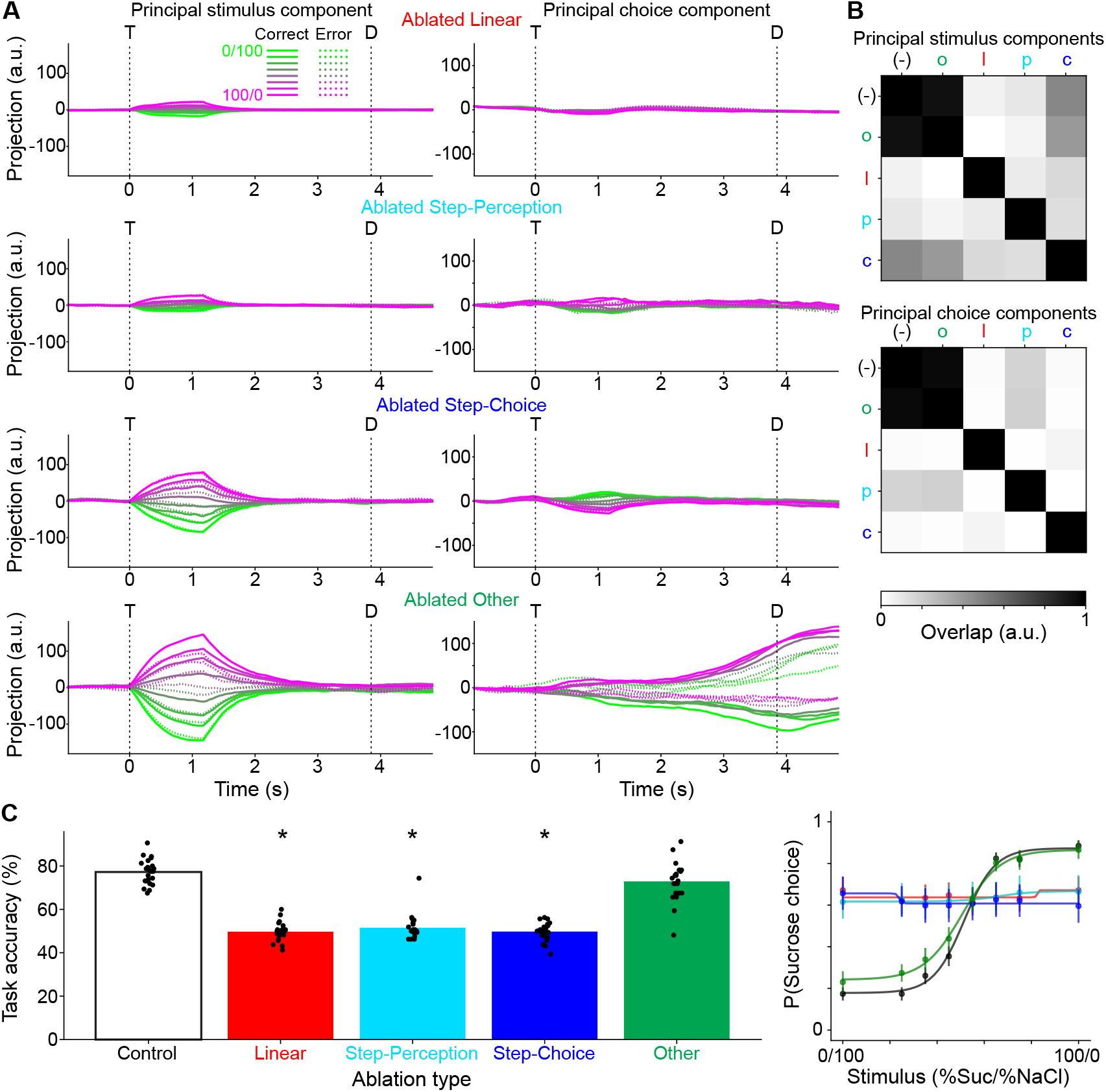
Effect of selective ablations on model dynamics and behavior. **A**: Model dynamics after selectively ablating linear coding units, step-perception coding units, step-choice coding units, or “other” units. Post-ablation pseudo-population activity is projected onto the stimulus-(left column) and choice-coding (right column) components identified in the control condition (i.e., the same ones in **Figure 6A-B**). Color scale corresponds to different mixture stimuli (%Sucrose/%NaCl); solid and dashed lines correspond to correct and error trials. T: time of stimulus onset; D: decision time. **B**: Pairwise overlaps between stimulus-(top) and choice-coding (bottom) components for control (-) and each ablation condition (o: “other,” l: linear, p: step-perception, c: step-choice). **C**: Behavioral performance of the model after selectively ablating categories of coding units. Left: across-model distributions of task accuracy vs ablation condition. Bars represent means. * indicates significant difference vs control condition (post-hoc paired t-test Bonferroni-adjusted *p* < 0.01). Right: psychometric functions fit to across-model mean probability of sucrose choice for different ablation conditions. Circles and error bars represent mean and s.e.m.

In terms of behavioral impact, all three coding types were also necessary for normal model performance, as task accuracy dropped significantly upon ablating any of them, whereas task performance was unaffected by ablating the “other” units (post-hoc paired t-tests with Bonferroni correction; control vs linear, *p* < 0.001; control vs step-perception, *p* < 0.001; control vs step-choice, *p* < 0.001; control vs “other,” *p* = 0.197; **Figure 7C**). Furthermore, the psychometric functions were not significantly different between the control and ablated “other” conditions, while they were different between the control and all other conditions (extra-sum-of-squares F-tests with Bonferroni correction; control vs linear, *p* < 0.001; control vs perception, *p* < 0.001; control vs choice, *p* < 0.001; control vs “other,” *p* = 0.235).

To determine the relative contributions of the constrained and unconstrained units to these results, we also carried out the coding type ablation simulations while restricting the ablation to the constrained or unconstrained sub-populations. The fractions of each coding category comprised of constrained units were 194/885 for linear, 147/987 for step-perception, 129/943 for step-choice, and 353/2092 for “other” units (**Table S4**). In terms of dynamics, there was a significant main effect of the constrained vs unconstrained population on stimulus coding activity (2-way within-subjects ANOVA with factors coding type [linear/step-perception/step-choice/other] and constraint [constrained/unconstrained]; constraint main effect *p* < 0.001; **Figure S7A**) and choice coding activity (2-way within-subjects ANOVA with factors coding type [linear/step-perception/step-choice/other] and constraint [constrained/unconstrained]; constraint main effect *p* < 0.001; **Figure S7B**); on average, ablating the unconstrained population impaired dynamics more, most likely due to its larger population size. Stimulus coding activity was sufficiently diminished by ablating linear or step-perception units restricted to the unconstrained population, but this was not the case for step-choice units at our *α* = 0.01 significance threshold (Dunnett test vs control; constrained linear, *p* = 0.291; unconstrained linear, *p* < 0.001; constrained step-perception, *p* = 0.661; unconstrained step-perception, *p* < 0.001; constrained step-choice, *p* = 0.953; unconstrained step-choice, *p* = 0.045; constrained “other,” *p* > 0.999; unconstrained “other,” *p* = 0.739; **Figure S7A**). Choice coding activity was impaired by ablating linear, step-perception, or step-choice units restricted to either the constrained or unconstrained populations (Dunnett test vs control; constrained linear, *p* < 0.001; unconstrained linear, *p* < 0.001; constrained step-perception, *p* < 0.001; unconstrained step-perception, *p* < 0.001; constrained step-choice, *p* < 0.001; unconstrained step-choice, *p* < 0.001; constrained “other,” *p* = 0.499; unconstrained “other,” *p* = 0.357; **Figure S7B**). In terms of behavioral performance, again there was a significant main effect of the constrained vs unconstrained population (2-way within-subjects ANOVA with factors coding type [linear/step-perception/step-choice/other] and constraint [constrained/unconstrained]; constraint main effect *p* < 0.001; **Figure S7C**), with unconstrained ablations having larger impact, and task accuracy was still reduced by ablating linear, step-perception, or step-choice units, regardless of their restriction to the constrained or unconstrained populations (Dunnett test vs control; constrained linear, *p* < 0.001; unconstrained linear, *p* < 0.001; constrained step-perception, *p* < 0.001; unconstrained step-perception, *p* < 0.001; constrained choice, *p* = 0.002; unconstrained choice, *p* < 0.001; constrained “other,” *p* = 0.649; unconstrained “other,” *p* > 0.999; **Figure S7C**).

Finally, to analyze the temporal aspect of coding unit relevance to dynamics and behavior, we conducted additional ablation simulations with refined targets and time windows, based on two periods of interest: the beginning, i.e., the first 1.2 s after stimulus onset, and the end, i.e., the last 1.2 s prior to decision. For each period, we silenced the units with a particular coding type label at some point within that period for the entirety of the period. In terms of dynamics, stimulus coding was significantly reduced only by silencing the linear units in the beginning period (Dunnett test vs control; linear beginning, *p* < 0.001; linear end, *p* = 0.990; step-perception beginning, *p* = 0.016; step-perception end, *p* = 0.999; step-choice beginning, *p* > 0.999; choice end, *p* = 0.994; **Figure S8A**), whereas choice coding was blunted by silencing step-choice units in the end period or by silencing linear units in either the beginning or the end periods (Dunnett test vs control; linear beginning, *p* < 0.001; linear end, *p* < 0.001; step-perception beginning, *p* = 0.508; step-perception end, *p* = 0.515; step-choice beginning, *p* = 0.969; step-choice end, *p* < 0.001; **Figure S8B**). In terms of behavioral performance, task accuracy was impaired by silencing linear units in the beginning period, step-perception units in the end period, or step-choice units in the end period (Dunnett test vs control; linear beginning, *p* < 0.001; linear end, *p* = 0.433; step-perception beginning, *p* = 0.242; step-perception end, *p* < 0.001; step-choice beginning, *p* > 0.999; step-choice end, *p* < 0.001; **Figure S8C**). Overall, this suggests dynamics and behavior are mostly driven by linear units in the beginning period and categorical units in the end period of the trial.

In summary, the effect of these ablations on dynamics and behavior suggests that the model was quite robust to perturbation in general, but sensitive to manipulations that targeted linear and step coding units.

## Discussion

GC plays a fundamental role in representing multiple sensory, affective, cognitive, and motor processes associated with a gustatory experience (***Yamamoto et al., 1985; Samuelsen et al., 2012; Gardner and Fontanini, 2014; Vincis and Fontanini, 2016***). This function is performed through time-varying patterns of neural activity that sequentially encode for different variables associated with a taste-related task. Neural dynamics in GC have been extensively studied both in single neurons and at the population level (***Sadacca et al., 2016; Mahmood et al., 2023; Mazzucato et al., 2015; Mendoza et al., 2024; Livneh and Andermann, 2021***), and their role in producing behavior is beginning to be elucidated (***Kusumoto-Yoshida et al., 2015; Mukherjee et al., 2019; Vincis et al., 2020***). Yet, the relationship among single neuron firing patterns, population dynamics, and behavior is not completely understood. Here we investigate how sub-populations of GC neurons defined by their single unit response profiles influence population dynamics and behavioral performance in the context of a taste mixture-based 2AC task.

By recording neuronal activity with high-density probes in the GC of behaving mice, we unveiled population and single neuron dynamics associated with a taste mixture 2AC task. We found that both population and single neuron activities go through a phase in which mixtures are linearly coded by their components’ concentration to a phase where stimuli are binarily coded. Additional analyses of single neuron activity show that units with binary coding could further be divided as either representing the predominant mixture component—that is, its overall taste quality (sweet vs salty, “step-perception”)—or the directional licking decisions cued by the stimulus (left vs right, “step-choice”). While some neurons showed only one coding pattern, others could have different response profiles at different times during the trial. Overall, the neurons whose tuning curves could be significantly fit by a linear or step function at any point in time were less than 50% of the total number of recorded neurons, with each specific type constituting no more than 27%.

To study the functional role of such groups of single units, we built RNN models (***Valente et al., 2022; Cohen et al., 2020***) of GC (one per session) and trained them to perform the taste mixture 2AC task while also reflecting the experimentally observed single neuron firing. The RNNs matched experimentally observed single neuron PSTHs, GC population dynamics, and rodent behavioral performance. Perturbing the model by removing different groups of neurons with distinct coding properties showed that linear and step coding neurons are necessary for neural population dynamics as well as behavioral performance. Ablation of the neurons that did not fit into the above categories had no impact on dynamics and performance, highlighting the functional importance of the single neuron firing patterns identified in this study.

Altogether the results presented in this study explain the role of single neuron firing patterns in a decision-making task and validate a data-driven, machine learning-based approach to modeling and generating hypotheses about the functional significance of system components (***Song et al., 2016; Barak, 2017; Yang and Wang, 2020; Valente et al., 2022***).

### Coding of task-related variables in GC of mice engaged in a taste mixture 2AC task

Since the pioneering work of ***Katz et al. (2001)***, it has been known that GC population dynamics can sequentially encode different aspects of a gustatory experience, from somatosensation to chemosensation to hedonic evaluation. More recent work has extended these findings to include GC population-level dynamics coding for taste-predictive cues, expectation, licking preparation, and abstract decision-making in the context of taste-based behavioral tasks (***Stapleton, 2007; Samuelsen et al., 2012; Livneh et al., 2017; Fonseca et al., 2018; Vincis et al., 2020; Lang et al., 2023***). For instance, ***Vincis et al. (2020)*** showed population activity trajectories separating according to sensory quality (sweet vs bitter) before licking decision (lick left vs lick right) in GC of mice performing a taste-based 2AC task.

The population-level analyses presented here add to the growing body of evidence for GC dynamics’ involvement in taste-based decision-making (***Miller and Katz, 2010; Fonseca et al., 2018; Vincis et al., 2020; Lang et al., 2023; Jezzini and Padoa-Schioppa, 2024; Kogan and Fontanini, 2024; Zheng et al., 2025***). In the context of the taste mixture-based 2AC task presented here, we found neurons that discriminate between predominantly-sucrose and predominantly-NaCl mixtures throughout the trial time course, with population firing rates featuring two peaks, one related to the sampling of taste and one preceding lateral licking. Decoding of population activity showed that the representation of taste mixtures peaks early while choice-related coding peaks just before lateral licking. We applied dimensionality reduction techniques, t-SNE and demixed PCA (***Kobak et al., 2016***), to extract the dominant trajectories coding for the task components in a neural space. The low-dimensional activity trajectories revealed a graded, linear separation based on mixture stimuli early on with sampling, and a binary separation based on selected choice later, prior to the decision. These population-level results, along with previous studies (***Kogan and Fontanini, 2024; Maier and Katz, 2013***), led us to search for single neurons with linear and step coding responses. Indeed, a subset of single units whose tuning curves represent mixture stimuli as a linear or step function were identified. Based on the analysis of correct and error trials, the step coding units could be further divided as either representing the predominant mixture component (step-perception) or the directional licking decisions cued by the stimulus (step-choice). It is worth mentioning that while some neurons displayed only one coding pattern, others showed coding patterns that could vary in time, providing evidence for multiplexing at the single neuron level. Regardless, the time course of the prevalence of linear and step-choice responses was consistent with the dynamics observed at the population level, with the former peaking during sampling and the latter before lateral licking. While the existence of these coding types suggests a role in driving population dynamics, the relatively limited presence of specific coding types in any time bin may cast doubts on their functional significance. In particular, step-perception units appeared in a uniformly low proportion throughout most of the trial and, surprisingly, peaked after the decision, perhaps serving as a feedback signal for learning by encoding the percept to be compared with the choice and outcome that just occurred. In this way, the percept could guide future adjustments of the choice following errors. The rarity of step-perception units and their post-decision peak therefore raised additional questions about the functional role of single unit coding patterns.

To address these questions on the functional role of the single unit activity described above, we relied on a modeling approach that would allow us to reproduce the experimentally recorded single unit activity as well as the behavioral performance for each session.

### Using RNNs to investigate the role of single neuron coding patterns

Determining the functional role of a particular brain region’s dynamics typically relies on optogenetic manipulations, which have become the gold standard due to their high temporal precision and the availability of specific genetically- and/or anatomically-targeted viral constructs (***Li et al., 2019; Emiliani et al., 2022***). Optogenetic silencing of GC and its inputs at different times during a trial has revealed the dynamic role of GC in encoding of palatability information (***Lin et al., 2021***), gaping behaviors (***Mukherjee et al., 2019***), expectation (***Kusumoto-Yoshida et al., 2015***), and taste-based decision-making (***Vincis et al., 2020***). As powerful as the approach is, it cannot be applied to selectively perturb neurons that are characterized by a specific coding pattern with no known genetic or connectivity signature. In other words, the specific coding populations we explored here cannot be manipulated by traditional optogenetic techniques. To address this gap, we relied on simulated manipulations in computational models where there is full control over all individual units.

Previous modeling efforts on GC have largely focused on replicating population-level phenomena (e.g., sequences of metastable states) starting from *a priori* architectures and tuning parameters by hand to reproduce population dynamics and behavioral performance (***Miller and Katz, 2010; Mazzucato et al., 2015, 2016, 2019; Lang et al., 2023***). While advancing our knowledge of the properties and origin of GC population dynamics, these efforts have not directly furthered our understanding of the role of single unit firing patterns. Here we turned to RNNs as an unbiased method for reproducing single unit firing activity (***Barak, 2017; Yang and Wang, 2020***). In particular, RNNs do not assume a specific synaptic matrix ahead of training. We took advantage of the relative ease of training RNNs (***Song et al., 2016; Paszke et al., 2019***) to learn single neuron activity and behavioral performance, capturing population dynamics in the process (***Perich et al., 2020; Cohen et al., 2020; Valente et al., 2022; Rajan et al., 2016***). Our RNN was composed of “constrained” and “unconstrained” units. Each constrained unit was paired with a target neuron from the experimental dataset, and the mismatch between model and experimental PSTHs was included in the loss function. Unconstrained units were included to represent unobserved neurons in the experimental recordings, and to add degrees of freedom to the network tasked with matching the constrained units’ activities to targets. Each unit was driven by stimulus input, recurrent input, and noise input. The external decision unit was trained to select the correct choice given the stimulus; thus, we modeled an ideal scenario in which stimuli have matched intensities and no motor biases exist. Unlike previous models (***Lang et al., 2023; Cisek et al., 2009***), an external preparatory input prior to decision was not necessary. While this is not proof that GC lacks preparatory inputs, it demonstrates that these dynamics can be produced internally.

The tractability of training RNNs via automatic differentiation typically comes at the expense of realism and mechanistic understanding when considering the network as a model of the brain. In contrast, spiking neural networks offer much increased biophysical plausibility, but are much harder to train (***Bohte, 2011; Li et al., 2021; DePasquale et al., 2016***) and often require *a priori* decisions about their architecture. Here we took additional measures to alleviate the trade-off between these two approaches by training a subset of neurons to reproduce the experimental PSTHs. This enhances biological realism and allows for RNN predictions to have more meaningful interpretations (***Cohen et al., 2020; Rajan et al., 2016***). For our ablation studies, we identified the sub-populations of coding units based on their response patterns in the simulations. We then ran new simulations while clamping the firing rates of units in specific sub-populations to 0. We found that all three coding types—linear, step-perception, and step-choice—were required for normal population dynamics and behavior. We showed the impact of these unit types on population-level trajectories measured with demixed PCA and on behavioral performance measured with a psychometric function. The removal of linear coding units made the network unable to produce appropriate stimulus-related and choice-related population activity as well as flattened the psychometric function. Surprisingly, the same effect was obtained by the ablation of step-perception units. Ablation of step-choice neurons had a less dramatic effect on stimulus-related population activity but flattened choice-related trajectories. Crucially, ablating all other neurons (which were much more plentiful) left dynamics and behavior intact. This lack of effect from ablation was not because the activity of those neurons was unrelated to the task. Indeed, many represented stimuli and upcoming choices, only in ways outside of our defined coding types. This demonstrates the disproportionate importance of linear, step-perception, and step-choice coding types to the model compared to alternative coding strategies, and highlights their relevance. Interestingly, our results emphasize the importance of step-perception neurons, which appear to be responsible for bridging early stimulus-evoked dynamics with successive choice-related activity.

The work presented here leaves some open questions that present promising avenues for future research. Accumulating evidence indicates that GC’s ongoing, taste-evoked, and decision-making activity is supported by metastable dynamics (***Jones et al., 2007; Mazzucato et al., 2015; Sadacca et al., 2016; Lang et al., 2023***), activity manifested at the ensemble level as coordinated changes between internal (hidden) states. Although this work focuses on single-unit activity, it does so in the context of population dynamics. We therefore believe that our framework is compatible with alternative approaches centered on ensemble activity. For instance, neurons with the coding patterns identified here could coordinate with others to participate in the formation of different metastable states. However, our model was not specifically constructed to exhibit metastability as we fit it to single neuron activities and behavioral performance without enforcing the structural connectivity constraints required to produce metastable dynamics (***Mazzucato et al., 2015***). Presumably, though, the most faithful model of GC would capture both single unit coding types and metastability, and future work could explore the implementation of a constrained fitting procedure that achieves this. Recent evidence also indicates that discrimination learning on this mixture task (that is, learning to improve difficult mixture pair discriminations via additional training) is associated with an increase in choice selective cells during the later delay period (***Kogan and Fontanini, 2024***). The mechanistic origins of this finding are unclear, and a modified version of the model presented here may prove to be a useful tool for clarifying them.

## Conclusions

In conclusion, our work shows that GC, a well-studied model for understanding cortical dynamics in sensory areas, can encode taste mixtures linearly or categorically, and choices categorically, during a mixture-based decision-making task. These phenomena are observed at the population level as well as at the single neuron level, and our findings are consistent with a dynamic progression of coding from representation of stimulus information to decision-making, with coding of perceptual category providing a bridging signal. The different types of coding sub-populations all make essential contributions to population dynamics and behavioral performance in our model of GC, underscoring the relevance of even small groups of neurons encoding task-relevant variables in very specific ways. It is worth noting that the neurons outside of these groups, whose activity was not necessary for normal population dynamics or behavioral performance in this particular task, may be critical for other taste-based decision-making tasks. Indeed, these “other” neurons may constitute a reservoir from which functionally-significant units emerge during learning according to task-specific demands. We believe the modeling approach taken here is a powerful means of analyzing the impact of single units dynamics on network activity and performance and makes a case for data-driven, interpretable RNNs as useful tools in neuroscience that compromise between realistic biophysical models and “black box” machine learning models.

While the research presented here focuses on GC, its implications go beyond taste. Both the findings and the approach are relevant for understanding population and single neuron dynamics in areas where sensory, cognitive, and motor activity are jointly encoded.

## Methods

### Stereotaxic surgeries

Mice were anesthetized using a cocktail of ketamine (70 mg/kg) and dexmedetomidine (1 mg/kg) via intraperitoneal injection. After the animal was fully anesthetized, the head was shaved and cleaned with iodine and 70% ethanol. The animal was then transferred onto a stereotaxic apparatus. During the surgery, the depth of anesthesia was monitored via visual inspection of breathing rate, toe pinch reflex, and whisking. A heating pad was used to maintain body temperature. After the skin was excised, the skull was exposed and cleaned with saline, dry swabs, and 70% ethanol. A small amount of Vetbond (3M) was used to secure the skin around the edge of the incision. A pencil was used to trace the coronal, interfrontal, sagittal, and lambdoid sutures. GC craniotomy sites (AP: +1.2 mm, ML: ±3.7 mm relative to Bregma) were marked with a permanent marker and covered with Kwik-Sil (World Precision Instruments). A craniotomy site above the cerebellum was drilled and a ground wire soldered to a male pin was placed (A-M system, Cat.No.786000). The midline of a custom head bar was aligned with the interfrontal and sagittal suture markings and positioned 1 mm posterior to Bregma. It was then secured with dental acrylic (C&B Metabond), covering both the skull and the top of the head bar. DV coordinates of Bregma and Lambda were remeasured, and a calibration point was marked on the head bar for stereotaxic reference.

### Immunohistochemistry

Mice were deeply anesthetized with 220 mg/kg pentobarbital sodium (390 mg/ml) and were perfused with phosphate buffer saline (PBS) followed by 4% paraformaldehyde (PFA) in PBS. After post-fixing overnight in 4% PFA, the brains were sliced at 50 µm with a vibratome (Leica VT-1000S). The brain slices were counterstained with Hoechst 33342 (1:5000 dilution, H3570, Thermo Fisher, Waltham, MA). Sections were mounted, cover-slipped, and imaged using a fluorescent microscope (Olympus BX51WI).

A Python-based GUI for Histological E-data Registration in Brain Space (HERBS) was used to register the slice images onto Allen CCF mouse atlas based on anatomical features for 2D and 3D visualization (***Fuglstad et al., 2023***; github.com/Whitlock-Group/HERBS). Reconstruction and visualization of electrode track trajectories was performed with open-source Allen CCF Tools (***Shamash et al., 2018***; github.com/cortex-lab/allenCCF) in a custom MATLAB script.

### Statistical tests

For simple distribution comparisons, we used Mann-Whitney U tests (i.e., rank-sum tests) or t-tests (Python: Scipy *stats*). For comparing proportions, we used Chi-squared tests or 2-tailed binomial tests (Python: Scipy *stats*). For within-subjects comparisons with more than two levels, we used 1-way repeated measures ANOVAs (Python: AnovaRM from *statsmodels*) and followed up significant results with post-hoc paired t-tests (Python: Scipy *stats*). Bonferroni corrections were implemented by multiplying *p* values by *K*-choose-2, where *K* is the number of levels. For within-subjects comparisons with two predictors, we used 2-way repeated measures ANOVAs (MATLAB: *fitrm* and *ranova*; MathWorks) to analyze interactions and main effects. To compare all groups to a single control group post hoc, we used Dunnett’s test (Python: Scipy *stats*). Additional statistical tests (such as extra-sum-of-squares F-test; see below) were implemented with custom code in Python or MATLAB. Significance level was taken as *α* = 0.01.

### Behavioral data

All psychometric curves (**Figures 1B, 5B, 7C**) are least-squares fitted 4-parameter logistic functions of the form:

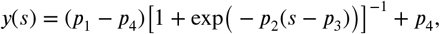

where *y* is the probability of a sucrose choice, *s* ∈ (0, 100) is the %Sucrose of the stimulus, *p*_1_ is the upper asymptote, *p*_2_ is the slope parameter, *p*_3_ is the inflection point, and *p*_4_ is the lower asymptote. Rather than averaging psychometric curves over sessions/models, a single psychometric was always fitted to the session-/model-averaged response as a function of the stimulus. Fitting was performed with Python’s *scipy* library, using the *curve_fit* method from the *optimize* module, with restrictions on the parameters: 0 ≤ *p*_1_ ≤ 1; 15 ≤ *p*_3_ ≤ 85; 0 ≤ *p*_4_ ≤ 1.

Psychometric curves were compared to each other using an extra-sum-of-squares F-test. The F statistic was calculated as (***Motulsky and Christopoulos, 2004; Maxwell et al., 2017***):

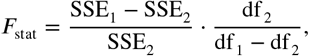

where SSE_1_ is the sum of squared errors when fitting all data points with a single curve, SSE_2_ is the sum of squared errors when fitting the separate data points with two separate curves, df _1_ = *N*_points_ − 4, and df _2_ = *N*_points_ − 8 (*N*_points_ is the total number of data points). The *p* value was calculated as the area under the F(df _1_ − df _2_, df _2_) distribution to the right of *F*_stat_ and evaluated at the *α* = 0.01 significance level. Bonferroni corrections were applied by multiplying *p* by *K*-choose-2, where *K* is the total number of psychometrics being considered.

### Electrophysiological data acquistion

Prior to each recording session, Neuropixel 1.0 probes were coated with Vybrant™ DiI (Thermo-Fischer) or neuro-DiO (Biotium). Before recording, the animal was placed in an induction chamber under 2.5% isoflurane for 2 to 3 minutes and then placed on a stereotaxic frame where anesthesia was maintained with 1 to 2% isoflurane. Dental acrylic over the Kwik-Sil was removed to expose the craniotomy site. A craniotomy was drilled based on the marker and cleaned with gel foam and saline. The animal was then transferred to the behavior platform. A multi-probe motorized manipulator system was used for recordings (New Scale Technologies). Bregma and Lambda were calibrated based on their relative distances from the calibration point. Neuropixels trajectory explorer with the Allen CCF mouse atlas was used to visualize the probe location in the brain (github.com/petersaj/neuropixels_trajectory_explorer). Craniotomies were kept moist with frequent application of saline during recordings and were sealed with Kwik-Sil and covered with a thin layer of dental cement after recording. The open-source software package SpikeGLX (billkarsh.github.io/SpikeGLX/) was used for data acquisition.

### Spike sorting

Spike sorting was performed with Kilosort 2 and Kilosort 4 (***Pachitariu et al., 2024***; github.com/MouseLand/Kilosort) and sorted clusters were manually curated using Phy (Cyrille Rossant, International Brain Laboratory) and custom MATLAB scripts. Units were identified with distinct clusters in waveform principal component space and a clear refractory period (> 1 ms) in auto-correlation histograms.

### Firing rate data

To analyze firing rate dynamics with respect to two discrete trial events—the time of the first central lick, T, and the time of the first lateral lick, D—simultaneously, we used a warped time scale. Spike trains were aligned to T, and a fixed number of bins (77) was used to calculate firing rates between T and D. This inter-event interval, IEI = D – T, varied from trial to trial, with an average duration of 3.85 s across the entire dataset (thus, the mean bin duration was ~50 ms). The fixed number of bins ensured firing rate timeseries could be aligned to both T and D across all trials from all sessions. Firing rates before T and after D were calculated using 50 ms bins. After averaging over trials to construct PSTHs, firing rate activities were smoothed using acausal Gaussian kernels 11 bins wide.

### auROC

Area under the receiver operating characteristic curve (auROC) was used to measure each neuron’s average difference in firing rate between %Sucrose < %NaCl and %Sucrose > %NaCl trials over time. Each ROC curve was constructed as Pr(*R*_1_ > *θ*) vs Pr(*R*_2_ > *θ*) for *R*_1_ the firing rate in %Sucrose < %NaCl trials, *R*_2_ the firing rate in %Sucrose > %NaCl trials, and *θ* the threshold parameter. Thus, an auROC value of 0 indicated firing rates in predominantly-sucrose trials were always greater, a value of 1 indicated firing rates in predominantly-NaCl trials were always greater, and a value of 0.5 represented complete overlap between distributions of firing rates in both trial types. Peak auROC values were identified as those with greatest absolute difference from 0.5, and neurons were labeled as sucrose- or NaCl-”preferring” based on whether this peak value was closer to 0 or 1.

### Responsivity and selectivity analyses

In line with previous work (***Kogan and Fontanini, 2024***), we classified neurons as responsive in the sampling (T to T + 0.5 s) or delay (D – 0.5 s to D) periods if their firing rate distributions were significantly different from baseline (T – 3 s to T – 2.5s for sampling baseline; D – 5.5 s to D – 5 s for delay baseline). We then checked responsive neurons to see if their firing rate distributions within sampling or delay periods were significantly different between predominantly-sucrose and predominantly-NaCl trials—if so, they were considered selective. All statistical comparisons were done via Mann-Whitney U tests at the *α* = 0.01 significance level. Only correct trials were used. We applied the same analyses to RNN units but with T – 0.5 s to T as the baseline for both sampling and delay periods.

### t-SNE

A t-distributed stochastic neighbor embedding (t-SNE) was used as a non-linear dimensionality reduction approach for visualizing pseudo-population firing rate trajectories. This method aims to preserve the true distance structure among points in the original, high-dimensional space when mapping to the low-dimensional embedding space. We concatenated correct trial PSTHs to construct an (*N*_stim_ · *N*_time_) × *N*_neu_ pseudo-population matrix for *N*_stim_ = 8 the number of unique stimuli, *N*_time_ = 117 the total number of time points, and *N*_neu_ = 626 the total number of neurons in the pseudo-population. The number of columns in the matrix was then reduced to 2 via the embedding—we used the *TSNE* class from the *manifold* module of Python’s *scikit-learn* library with default parameters.

### dPCA

A demixed principal component analysis (dPCA) was performed as a linear, supervised alternative to t-SNE for dimensionality reduction. Detailed methods are described in ***Kobak et al***. (***2016***). Briefly, this analysis began by organizing the firing rate data in a multi-dimensional array *X* of shape *N*_trials_ ×*N*_neurons_ ×*S* ×*Q*×*T*, where *T* = 117 is the number of time bins, *Q* = 2 is the number of choices, *S* = 8 is the number of stimuli, *N*_neurons_ = 626 is the total number of recorded neurons, and *N*_trials_ is the maximum number of trials across all sessions for any stimulus and choice combination (correct and error trials are included). Whenever a session had no trials for a particular stimulus/choice combination, we fit simple linear models to each time bin’s available data and used them to predict the missing data:

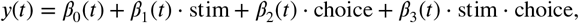

where stim (in %Sucrose) is the stimulus value, choice ∈ {0, 1} is the animal’s choice (0 for “NaCl choice” and 1 for “sucrose choice”), *y*(*t*) is the predicted firing rate in time bin *t*, and the *β*(*t*)s are the fitted regression weights. At least 2 trials per session and stimulus/choice combination are required for optimizing the regularization hyperparameter in dPCA (if only 1 existed, it was copied), though the main dPCA analysis operates on the trial-averaged data matrix 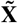 of shape *N*_neurons_ ×*SQT*. The analysis partitions this matrix into marginalized contributions from each variable, 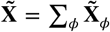, where *ϕ* ∈ {*s, q, t, sq, st, qt, sqt*} is a variable combination (*s* for stimulus, *q* for choice, *t* for time; we joined *s* with *st, q* with *qt*, and *sq* with *sqt*), then minimizes the loss *L*_*ϕ*_ for each:

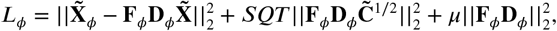

where **F**_*ϕ*_ and **D**_*ϕ*_ are encoders and decoders, respectively, 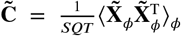 is the average covariance matrix over all parameter combinations (⟨·⟩_*ϕ*_ denotes averaging over *ϕ* and (·)^T^ denotes matrix/vector transposition), *μ* is the regularization hyperparameter, and ||·|| _2_ denotes the Frobenius norm for matrices and the 2-norm for vectors. After fitting, one-dimensional projections of pseudo-population activity were obtained by calculating the inner product of **d**_*ϕ*_ and the *N*_neurons_-dimensional vector over time, where **d**_*ϕ*_ is the row of **D**_*ϕ*_ that maximized projected variance.

Overlaps between components obtained via dPCA were calculated as an absolute cosine similarity between vectors:

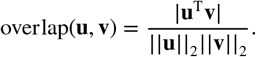

### Minimum-distance decoding

For each session, we trained minimum-distance classifiers to decode several task-relevant variables from population firing rate activity over time. These custom classifiers may be thought of as variants of 1-nearest neighbor classifiers where neighbors are centroids (class averages). At each point in time, all trials were represented as population firing rate vectors and their associated labels {(**r**_*i*_, *l*_*i*_)} for *i* indexing trials and *l*_*i*_ ∈ *U*, the set of unique labels (e.g., for decoding Choice, *U* = {Left, Right}). Each trial i was held out, and mean population firing rate vectors were calculated for each label *u* ∈ *U* :

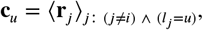

where ∧ denotes logical “and.”

The held-out trial was then assigned the class label corresponding to the nearest mean firing rate vector:

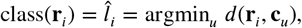

where *d* is a distance metric. We chose the Euclidean distance metric, *d*(**r**_*i*_, **c**_*u*_) = ||**r**_*i*_ − **c**_*u*_ ||_2_. The class-balanced leave-one-out test accuracy was calculated as:

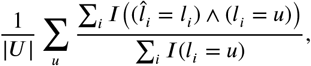

where |*U* | is the cardinality of *U*, i.e., the number of unique labels, and *I* is the indicator function:

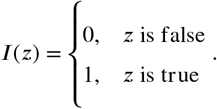

A one-tailed *α*-significance threshold for the session-averaged decoding accuracy was calculated from the binomial distribution as *k*/*N*, where *N* is the average number of trials and *k* is the minimum number of hits such that the tail probability to the right of *k* is still below *α*. That is:

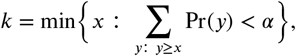

where Pr(*y*) is the probability mass function of a binomial random variable parameterized by *N* and *p*. We used *α* = 0.01, *N* = 137 (the average number of trials across all sessions), and *p* ∈ {1/8, 1/2} (the theoretical chance levels of decoding each variable).

We used a 2-way within-subjects ANOVA with factors decoding (stimulus or choice) and time (sampling or delay) to compare the time courses of decoding. Each decoding time course had its theoretical chance level subtracted prior to the test, and decoding time courses were averaged within the first 10 bins after T (T to ~T + 0.5 s) and last 10 bins before D (~D – 0.5 s to D) for sampling and delay, respectively.

### Response profiles

Single unit response profiles were analyzed with a least-squares regression-based pipeline similar to that of ***Maier and Katz (2013***). Let *x*_*k,t*_ represent a single neuron’s pre-processed firing rate data (i.e., already time-warped and smoothed) for *k* indexing trials and *t* indexing time. Let *σ*(*k*) be the stimulus administered on trial *k* (in terms of %Sucrose, for simplicity) and *o*(*k*) be the outcome of the trial (i.e., correct or error). For each neuron from each session, the response profile *r* was calculated as a function of the stimulus *s* (again, in terms of %Sucrose) for a given time window *w*:

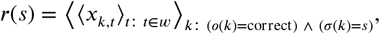

where again ⟨·⟩_*x*_ denotes averaging over *x* and ∧ denotes logical “and”. The shape of each response profile was then labeled “linear,” “step,” or “other” by comparing to template shapes: a 2-parameter line,

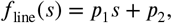

and a 3-parameter step function,

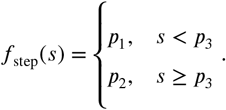

The comparison was carried out by finding the best fit (in the least-squares sense) for each template, and then comparing the resulting *F* values. The *F*_stat_ value for each template was calculated from the extra-sum-of-squares *F* formula above (**Methods: Behavioral data**), now applied to a fitted vs null model comparison rather than a nested model comparison (***Motulsky and Christopoulos, 2004; Maxwell et al., 2017***):

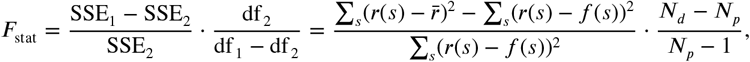

where SSE_1_ is the sum of squared error for the null model (the mean of the data), SSE_2_ is the sum of squared error for the fitted model, df _1_ and df _2_ are the corresponding degrees of freedom, 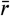 is the average of *r*(*s*), *N*_*d*_ is the number of data points, and *N*_*p*_ is the number of parameters. The shape of the response profile was then assigned as the template with the largest associated *F*_stat_ value, as long as this *F*_stat_ value’s corresponding *p* value (area under the F(df _1_ − df _2_, df _2_) distribution to the right of *F*_stat_) was less than *α* = 0.005 (0.01, Bonferroni-corrected for the number of tests). If this was not the case, the response profile’s shape was assigned as “other.” Least-squares fitting for *f*_line_ was performed as a constrained fit, subject to *f*_line_(*s*) ≥ 0 for all *s*, using Python’s *scipy* library (*minimize* method of the *optimize* module). Least-squares fitting for *f*_step_ was performed manually by varying *p*_3_ ∈ {40, 50, 60} and, for each, finding the optimal remaining parameters as 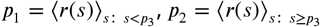.

To sub-classify neurons labeled “step” into “step-perception” and “step-choice,” we incorporated error trials. To be considered a “step-choice” neuron, two criteria had to be met: (i) the inflection point of the step response profile (*p*_3_) had to occur at *s* = 50; (ii) the average firing rate of the neuron had to be consistently (between correct and error trials) greater for trials where the same direction was chosen. For example, if the firing rate averaged over correct trials of *s* > 50 was greater than the average over correct trials of *s* < 50, then we required its firing rate averaged over error trials of *s* < 50 to be greater than the average over error trials of *s* > 50 (since that would indicate a consistent “preference” for trials of one lick direction). More explicitly, we calculated *δ*_*x*_ = ⟨*r*_*x*_(*s*)⟩_*s*: *s*>50_ −⟨*r*_*x*_(*s*)⟩_*s*: *s*<50_ for *x* ∈ {correct, error} (where *r*_correct_ (*s*) = *r*(*s*) and *r*_error_ (*s*) is obtained by replacing the condition *σ*(*k*) = correct with *σ*(*k*) = error in the above definition of *r*(*s*)) and required *δ*_correct_ *δ*_error_ < 0 for the “step-choice” label. If no error trials existed for stimulus values on either side of *s* = 50 (relevant for some simulated data), “step” neurons were not sub-classified.

We examined the proportions of all neurons pooled over all sessions and subjects that were classified as each shape at the beginning and end of the trial using windows *w*_1_ = [T, T + 0.5 s] and *w*_2_ = [D − 0.5 s, D] on the un-warped time scale. Chi-squared tests of proportions were done without Yates’ correction, using *chi2_contingency* from the *stats* module of Python’s *scipy* library. We also visualized the proportions over the full trial time course by using a moving window 4 bins wide (~200 ms) on the warped time scale, stepped by 4 bins (the final window was only 1 bin wide since the total number of time points was not divisible by 4).

### RNN: Components and dynamics

Recurrent neural network models were inspired by ***Cohen et al***. (***2020***) and adapted from opensource code (***Valente et al., 2022***; github.com/adrian-valente/lowrank_inference). Each RNN used here is comprised of *N*_c_ constrained and *N*_u_ unconstrained artificial units, for *N*_c_ equal to the number of simultaneously recorded neurons in the corresponding experimental session and the total number set at *N* = *N*_c_ + *N*_u_ = round(5.88 · *N*_c_). The additional decision unit, described later, is not included in the total count.

The internal activity of all units in the network, **h** ∈ ℝ^*N*^, is influenced by external stimulus input, noise input, and recurrent input. The external stimulus input, *m*(**x**), is the mixture stimulus modeled as a 2-dimensional vector—e.g., 25/75 (%Sucrose/%NaCl) is **x** = [0.25, 0.75]^T^—passed through the non-linear mapping *m* : ℝ^2^ → ℝ^*N*^ defined by *m*(**x**) = **A**^(2)^tanh(**A**^(1)^**x**) for matrices **A**^(1)^ ∈ ℝ^100×2^, **A**^(2)^ ∈ ℝ^*N*×100^, and tanh applied element-wise to vectors. The noise input is **η** for *η*_*i*_ ~ 𝒩 (0, *σ*_*η*_) resampled at each time step from a Gaussian distribution with mean 0 and standard deviation *σ*_*η*_ = 0.05/*α* (*α* defined below). The recurrent input is **W**_rec_*f* (**h** + **b**) for the synaptic matrix **W**_rec_ ∈ ℝ^*N*×*N*^, the input bias **b** ∈ ℝ^*N*^, and the transfer function *f*, applied element-wise to vectors, is a rectified linear function with a maximum value of 80, i.e., *f* (*z*) = min(max(*z*, 0), 80). The output of the transfer function is interpreted as a firing rate, i.e., **r** = *f* (**h** + **b**). Network activity evolves according to:

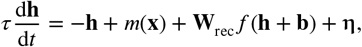

for time constant *τ*. We integrated this equation in discrete time using the forward Euler algorithm with step size *α* = d*t*/*τ* = 0.2. The model’s decision over time is governed by an additional decision unit that is functionally external from the network (i.e., it does not appear in the equation for **h** above). This unit is denoted by *z* and its internal activity is driven by the firing rates of all *N* units in the network:

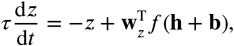

for weight vector **w**_*z*_ ∈ ℝ^*N*^. This equation is integrated in parallel with the one for **h** and the model’s binary choice is determined by interpreting *c* = tanh(*z*) as “sucrose choice” if *c* > 0 and as “NaCl choice” if *c* < 0.

### RNN: Training

The goal during training is for the model to respond to any particular stimulus input by producing the correct choice (i.e., value of *c*) during a pre-defined decision window as well as constrained firing rate activities (i.e., first *N*_c_ components of **r**) that match the correct trial PSTHs of the neurons in the corresponding session. Thus, we define an overall loss as:

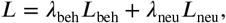

for behavioral loss *L*_beh_, neural loss *L*_neu_, and associated weights *λ*_beh_ = 150 and *λ*_neu_ = 1 to counter-balance the different error scales. The behavioral loss is:

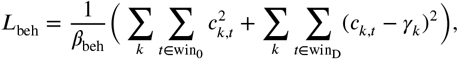

where *k* indexes stimuli, *t* indexes time, win_0_ = [T − 1 s, T] is the pre-stimulus window for T the stimulus delivery time, win_D_ = [D − 0.1 s, D] is the decision window for D the decision time, *c*_*k,t*_ is the network’s value of *c* in response to stimulus *k* at time *t, γ*_*k*_ is the correct decision in response to stimulus *k* (e.g., for *k* = 0/100, *γ*_*k*_ = −1; for *k* = 100/0, *γ*_*k*_ = +1), and *β*_beh_ is the total number of terms in the sums. The neural loss is:

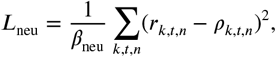

where *n* indexes constrained neurons, *r*_*k,t,n*_ is the network’s firing rate output for neuron *n* in response to stimulus *k* at time *t, ρ* is the experimentally observed correct trial PSTHs, and *β*_neu_ is the total number of terms in the sum.

Training was carried out in PyTorch (***Paszke et al., 2019***): *L* was minimized with respect to the network’s trainable parameters—**A**^(1)^, **A**^(2)^, **W**_rec_, **b, w**_*z*_, and the initial value of **h**—via backpropagation, using the Adam optimizer with a learning rate of 0.01 and gradient clipping above 1.0. Elements of **A**^(1)^ and **A**^(2)^ were randomly initialized as 𝒩 (0, 1). Elements of **W**_rec_ were randomly initialized as 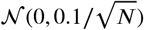, with self-connections prohibited. Elements of **w**_*z*_ were randomly initialized as 𝒩 (0, 1/*N*). Elements of **b** and the initial values of **h** were initialized at 0. Training proceeds for 2000 iterations or until *L* < 1.

### RNN: Simulations

Single trial simulations (both during and after training) were conducted for 5.9 s each, matching the time-warped window of analysis used for experimental data. The stimulus **x** was “on” 1 s after trial start and lasted for 1.2 s. When off, **x** = [0, 0]^T^. After training, simulations were carried out with persistent additional noise, changing the external input current to *m*(**x** + **ϵ**) with *ϵ*_*i*_ ~ 𝒩 (0, *σ*) resampled at each time step. The level of external noise, *σ*, was tailored to each model such that its overall task accuracy was within 5% of the animal’s accuracy from the corresponding experimental session (*σ* ranged from 0.10 to 1.15). We simulated 20 trials per stimulus, and choices were obtained from the sign of the average value of *c* over the decision window, which covers 0.1 s before the decision time (which occurs 3.9 s after stimulus start) up to the decision time.

For ablation simulations, elements of **r** = *f* (**h** + **b**) corresponding to firing rates of units belonging to specific sub-populations of interest (identified based on results from simulations without ablation) were clamped to 0 for all time, except in **Figure S8**, where the firing rates were clamped to 0 only in specific temporal windows. Ablations can result in the model becoming highly biased toward one choice direction; to limit inclusion of inferred missing data for the dPCA of model dynamics across ablation conditions, we excluded models that did not have at least 2 trials for each chosen direction under all conditions. Out of 23 models, the number of models passing criterion depended on the conditions considered and were 14 for **Figures 6A-B, 7A-B,S5A-B**, and **S6A-B**; 13 for **Figure S8A-B**; and 8 for **Figure S7A-B**.

## Acknowledgements

Work supported by R01DC018227 from NIH/NIDCD (A.F.), 1UF1NS115779 from NIH/NINDS Brain Initiative (AF and GLC), American Association of University Women (AAUW) International Fellowship (C.Y.Z), K12GM102778 from NIH/NGM (J.M.B).

---

**Supplementary Table 1.**
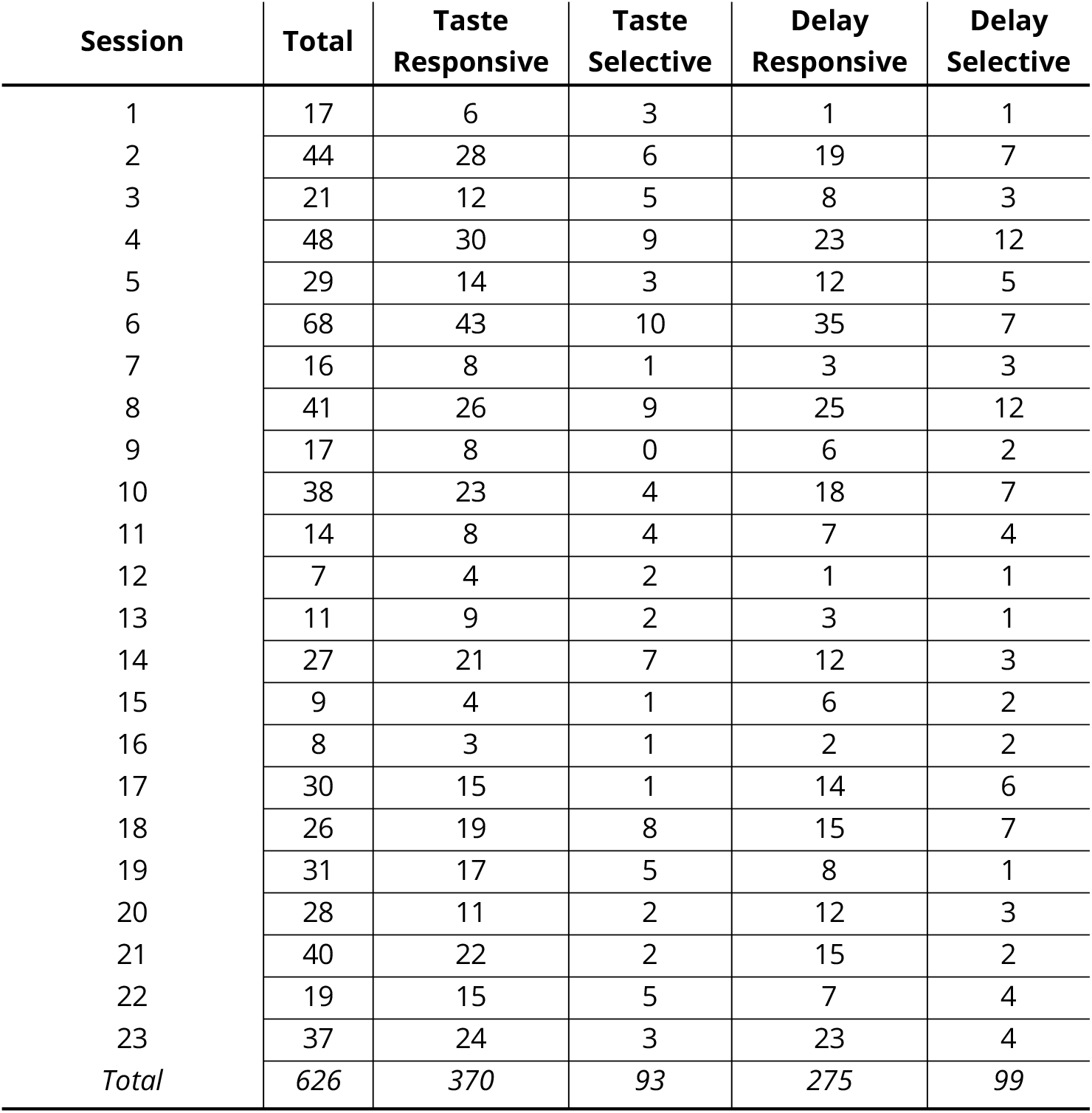
Session-by-session responsive and selective neuron counts for experimental data. Responsivity indicates a difference in firing rate distributions between baseline and a window of interest (from the first central lick to 500 ms after it for taste; from 500 ms before the first lateral lick to the first lateral lick for delay). Selectivity indicates a difference in firing rate distributions between categories within the window of interest (predominantly-sucrose vs predominantly-NaCl for taste; left vs right for delay).

**Supplementary Table 2.** Session-by-session neuron coding type counts for experimental data. Neurons are assigned coding type labels if they exhibit the response profile pattern in any time bin (as per analysis in **Figure 4C**) and, thus, the labels are not mutually exclusive.

| Session | Total | Linear | Step-Perception | Step-Choice | Other |
| --- | --- | --- | --- | --- | --- |
| 1 | 17 | 4 | 6 | 1 | 8 |
| 2 | 44 | 13 | 9 | 8 | 27 |
| 3 | 21 | 3 | 11 | 1 | 9 |
| 4 | 48 | 22 | 11 | 9 | 23 |
| 5 | 29 | 8 | 5 | 6 | 18 |
| 6 | 68 | 16 | 16 | 10 | 38 |
| 7 | 16 | 3 | 6 | 2 | 9 |
| 8 | 41 | 8 | 12 | 17 | 17 |
| 9 | 17 | 1 | 2 | 4 | 10 |
| 10 | 38 | 6 | 4 | 12 | 22 |
| 11 | 14 | 6 | 4 | 2 | 4 |
| 12 | 7 | 3 | 1 | 1 | 4 |
| 13 | 11 | 4 | 2 | 0 | 5 |
| 14 | 27 | 5 | 8 | 5 | 14 |
| 15 | 9 | 4 | 0 | 0 | 5 |
| 16 | 8 | 3 | 2 | 2 | 4 |
| 17 | 30 | 6 | 9 | 4 | 17 |
| 18 | 26 | 8 | 14 | 12 | 7 |
| 19 | 31 | 5 | 11 | 8 | 14 |
| 20 | 28 | 6 | 7 | 2 | 16 |
| 21 | 40 | 7 | 6 | 4 | 25 |
| 22 | 19 | 6 | 9 | 2 | 5 |
| 23 | 37 | 8 | 12 | 2 | 21 |
| *Total* | *626* | *155* | *167* | *114* | *322* |

**Supplementary Table 3.** Session-by-session responsive and selective unit counts for model data. Responsivity indicates a difference in firing rate distributions between baseline and a window of interest (from stimulus onset to 500 ms after it for taste; from 500 ms before the decision to the decision for delay). Selectivity indicates a difference in firing rate distributions between categories within the window of interest (predominantly-sucrose vs predominantly-NaCl for taste; left vs right for delay).

| Session | Total |  | Taste Responsive |  | Taste Selective |  | Delay Responsive |  | Delay Selective |  |
| --- | --- | --- | --- | --- | --- | --- | --- | --- | --- | --- |
|  | Con. | Unc. | Con. | Unc. | Con. | Unc. | Con. | Unc. | Con. | Unc. |
| 1 | 17 | 83 | 10 | 55 | 10 | 42 | 8 | 48 | 5 | 38 |
| 2 | 44 | 215 | 9 | 25 | 5 | 15 | 10 | 14 | 7 | 10 |
| 3 | 21 | 102 | 11 | 62 | 8 | 38 | 13 | 51 | 8 | 42 |
| 4 | 48 | 234 | 10 | 79 | 9 | 52 | 9 | 59 | 10 | 49 |
| 5 | 29 | 142 | 20 | 97 | 8 | 77 | 19 | 74 | 13 | 79 |
| 6 | 68 | 332 | 25 | 144 | 20 | 109 | 30 | 115 | 18 | 115 |
| 7 | 16 | 78 | 10 | 45 | 7 | 27 | 12 | 36 | 13 | 42 |
| 8 | 41 | 200 | 19 | 79 | 13 | 64 | 17 | 62 | 10 | 21 |
| 9 | 17 | 83 | 8 | 34 | 3 | 19 | 6 | 29 | 4 | 13 |
| 10 | 38 | 185 | 10 | 75 | 5 | 35 | 15 | 67 | 10 | 29 |
| 11 | 14 | 68 | 6 | 42 | 3 | 31 | 9 | 37 | 9 | 40 |
| 12 | 7 | 34 | 3 | 23 | 2 | 18 | 5 | 22 | 3 | 24 |
| 13 | 11 | 54 | 8 | 39 | 5 | 31 | 2 | 25 | 6 | 34 |
| 14 | 27 | 132 | 13 | 75 | 8 | 56 | 16 | 98 | 10 | 71 |
| 15 | 9 | 44 | 6 | 34 | 8 | 38 | 9 | 29 | 3 | 38 |
| 16 | 8 | 39 | 7 | 29 | 4 | 25 | 6 | 26 | 4 | 27 |
| 17 | 30 | 146 | 16 | 86 | 11 | 60 | 10 | 55 | 7 | 49 |
| 18 | 26 | 127 | 19 | 74 | 10 | 46 | 17 | 55 | 14 | 37 |
| 19 | 31 | 151 | 25 | 101 | 12 | 86 | 18 | 61 | 11 | 69 |
| 20 | 28 | 137 | 14 | 91 | 8 | 59 | 10 | 57 | 10 | 67 |
| 21 | 40 | 195 | 20 | 106 | 13 | 87 | 24 | 84 | 16 | 71 |
| 22 | 19 | 93 | 13 | 60 | 5 | 43 | 10 | 43 | 7 | 29 |
| 23 | 37 | 181 | 17 | 78 | 12 | 65 | 17 | 63 | 12 | 59 |
| Total | 626 | 3055 | 299 | 1533 | 189 | 1123 | 292 | 1210 | 210 | 1053 |

**Supplementary Table 4.** Session-by-session unit coding type counts for model data. Units are assigned coding type labels if they exhibit the response profile pattern in any time bin (as per analysis in **Figure 6C**) and, thus, the labels are not mutually exclusive. Con.: constrained units; Unc.: unconstrained units.

| Session | Total |  | Linear |  | Step-Perception |  | Step-Choice |  | Other |  |
| --- | --- | --- | --- | --- | --- | --- | --- | --- | --- | --- |
|  | Con. | Unc. | Con. | Unc. | Con. | Unc. | Con. | Unc. | Con. | Unc. |
| 1 | 17 | 83 | 5 | 17 | 6 | 32 | 6 | 42 | 10 | 28 |
| 2 | 44 | 215 | 8 | 9 | 5 | 18 | 9 | 14 | 33 | 191 |
| 3 | 21 | 102 | 6 | 13 | 4 | 29 | 2 | 36 | 14 | 56 |
| 4 | 48 | 234 | 9 | 37 | 8 | 46 | 3 | 25 | 37 | 175 |
| 5 | 29 | 142 | 11 | 57 | 12 | 61 | 3 | 57 | 11 | 52 |
| 6 | 68 | 332 | 13 | 53 | 8 | 70 | 11 | 80 | 47 | 204 |
| 7 | 16 | 78 | 11 | 17 | 12 | 34 | 9 | 32 | 3 | 39 |
| 8 | 41 | 200 | 12 | 21 | 4 | 28 | 1 | 4 | 27 | 163 |
| 9 | 17 | 83 | 4 | 12 | 7 | 23 | 3 | 9 | 10 | 56 |
| 10 | 38 | 185 | 12 | 28 | 3 | 19 | 8 | 27 | 26 | 137 |
| 11 | 14 | 68 | 7 | 33 | 8 | 40 | 4 | 32 | 4 | 21 |
| 12 | 7 | 34 | 2 | 15 | 2 | 22 | 0 | 19 | 3 | 6 |
| 13 | 11 | 54 | 4 | 32 | 2 | 38 | 4 | 35 | 4 | 10 |
| 14 | 27 | 132 | 10 | 39 | 9 | 50 | 8 | 44 | 14 | 52 |
| 15 | 9 | 44 | 3 | 19 | 0 | 3 | 4 | 39 | 4 | 4 |
| 16 | 8 | 39 | 4 | 15 | 5 | 17 | 3 | 29 | 0 | 6 |
| 17 | 30 | 146 | 4 | 22 | 4 | 38 | 7 | 46 | 20 | 88 |
| 18 | 26 | 127 | 16 | 31 | 17 | 39 | 5 | 27 | 6 | 67 |
| 19 | 31 | 151 | 14 | 38 | 6 | 50 | 14 | 58 | 12 | 63 |
| 20 | 28 | 137 | 9 | 53 | 3 | 49 | 4 | 41 | 18 | 62 |
| 21 | 40 | 195 | 16 | 48 | 10 | 62 | 11 | 57 | 19 | 102 |
| 22 | 19 | 93 | 5 | 31 | 4 | 25 | 6 | 28 | 9 | 48 |
| 23 | 37 | 181 | 9 | 51 | 8 | 47 | 4 | 33 | 22 | 109 |
| Total | 626 | 3055 | 194 | 691 | 147 | 840 | 129 | 814 | 353 | 1739 |

**Supplementary Figure 1.**
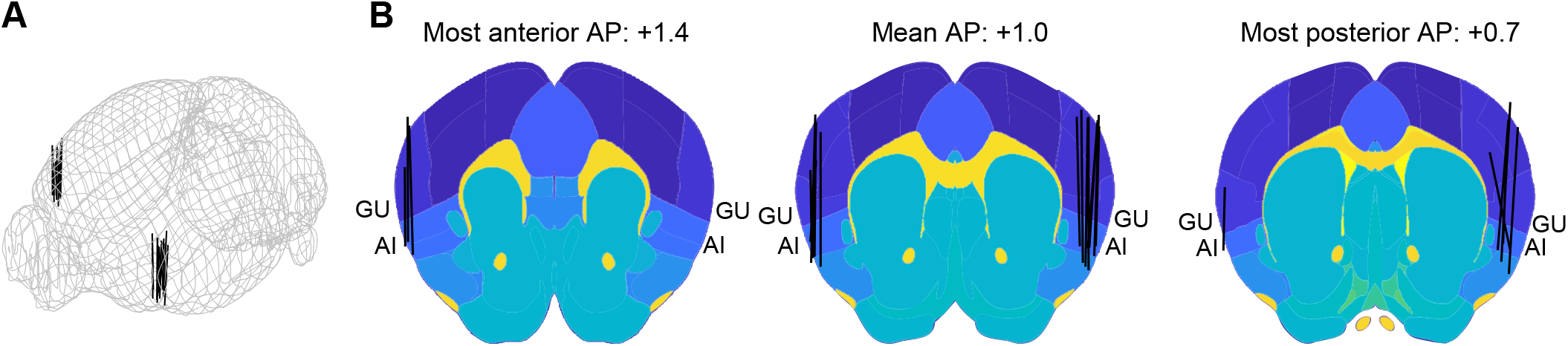
Neuropixels probe trajectory reconstruction. **A**: 3D reconstruction of the 23 probe trajectories from the experimental dataset. **B**: 2D reconstruction of the same 23 probe trajectories, overlaid on the Allen Brain Atlas at varying anteroposterior (AP) distances (relative to Bregma in mm) around GC. At these coordinates, both GU (gustatory areas) and AI (anterior insular areas) account for GC. Reconstructions performed with open-source Allen CCF Tools (***Shamash et al., 2018***; github.com/cortex-lab/allenCCF).

**Supplementary Figure 2.**
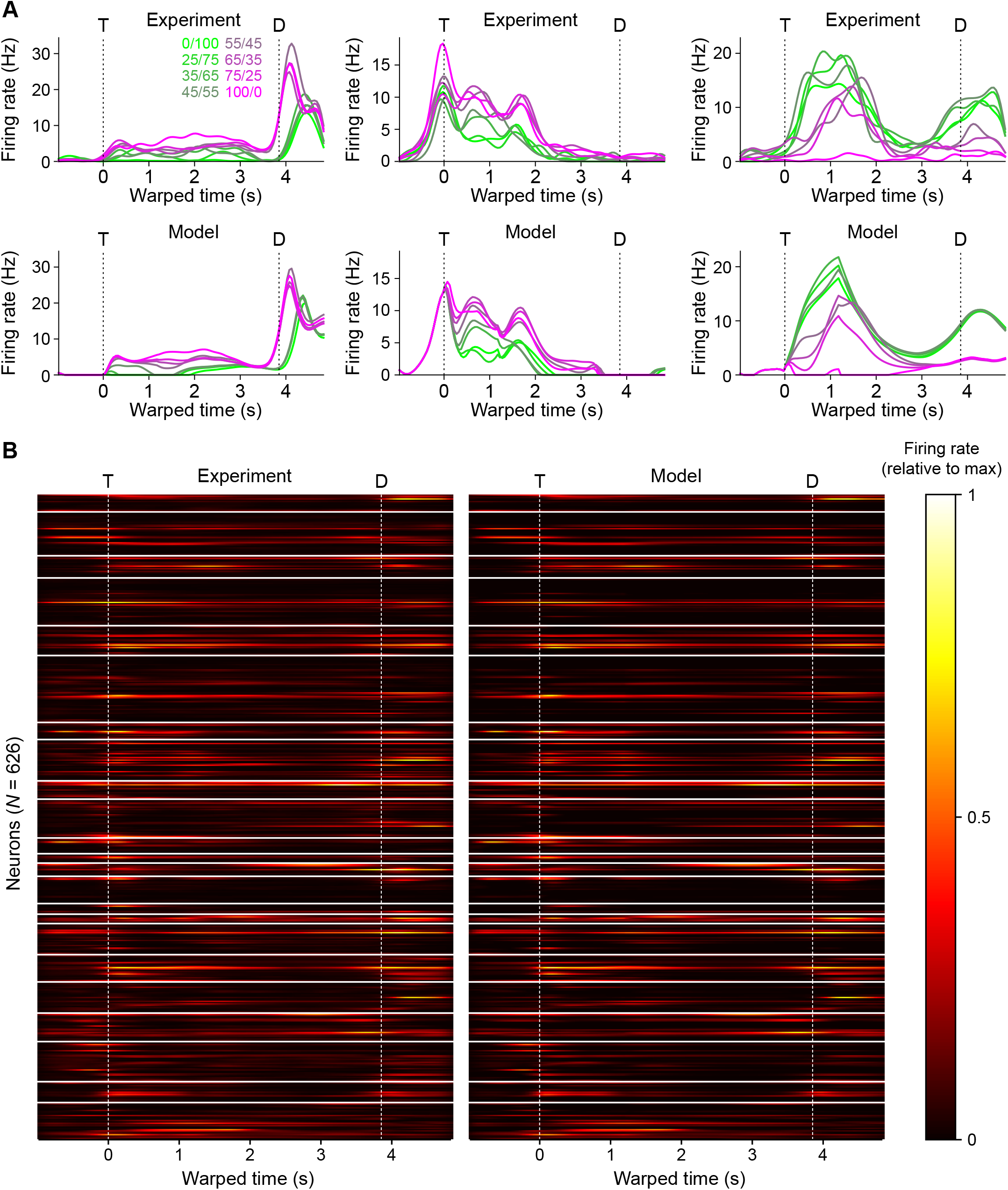
Model constrained unit activity. **A**: Three examples (columns) of experimentally-observed PSTHs (top) and corresponding model unit firing rate activities trained to match them (bottom). Color scale corresponds to different mixture stimuli (%Sucrose/%NaCl). **B**: Comparison of firing rate activities (stimulus-averaged PSTHs) between all experimental neurons and their corresponding model constrained units. Vertical whitespace separates individual sessions/models. Firing rates are normalized to the maximum within each session. T: time of first central lick/stimulus onset; D: time of first lateral lick/decision time.

**Supplementary Figure 3.**
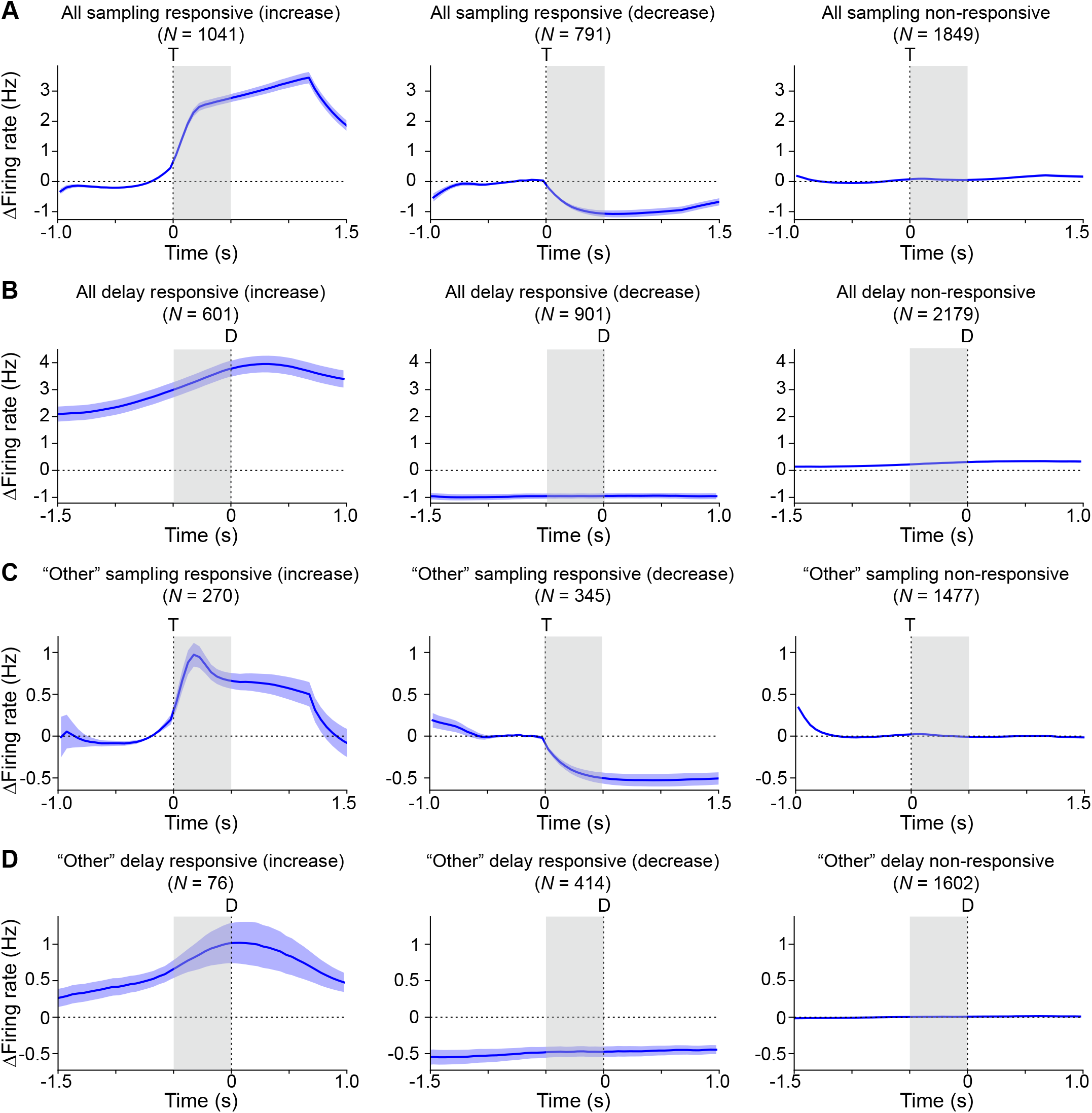
RNN unit responsiveness. **A**: Activity of all RNN units grouped by responsiveness during the sampling period. If the unit’s firing rate distribution during the sampling period (T to T + 0.5 s for T the stimulus onset time) was significantly different from its baseline (T – 0.5 s to T) firing rate distribution, it was sampling responsive and grouped by whether its mean firing rate increased (left) or decreased (middle); otherwise it was non-responsive (right). **B**: Activity of all RNN units grouped by responsiveness during the delay period. Same as **A** except the firing rate distribution of interest is calculated over D – 0.5 s to D for D the decision time. **C** and **D**: Same as **A** and **B**, respectively, except that the only units considered are those labeled “other” by the response profile analysis of **Figure 6C**. Firing rates are expressed relative to baseline, and traces are population mean ± s.e.m.

**Supplementary Figure 4.**
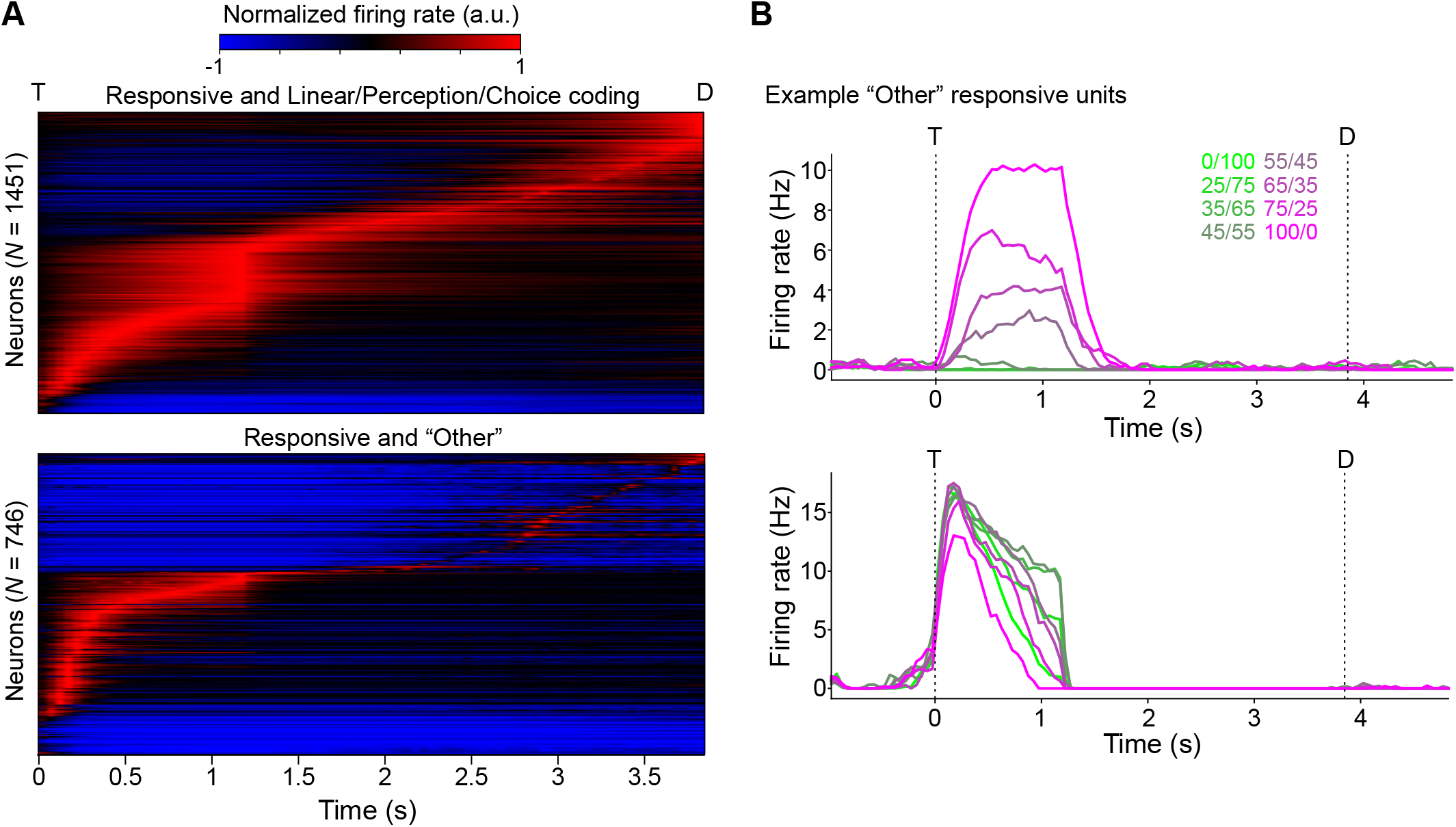
RNN unit activity patterns. **A**: Heatmaps of firing rate activities for units that responded significantly during the sampling and/or delay periods, broken down into coding units (linear, step-perception, and/or step-choice) (top) and “other” units (not linear, not step-perception, and not step-choice) (bottom). Firing rates are expressed relative to baseline and normalized to the maximum absolute value. T: time of stimulus onset; D: decision time. **B**: Two example “other” unit responses. Both respond significantly during the sampling period, but neither response pattern matches the linear or step templates. Color scale corresponds to different mixture stimuli (%Sucrose/%NaCl).

**Supplementary Figure 5.**
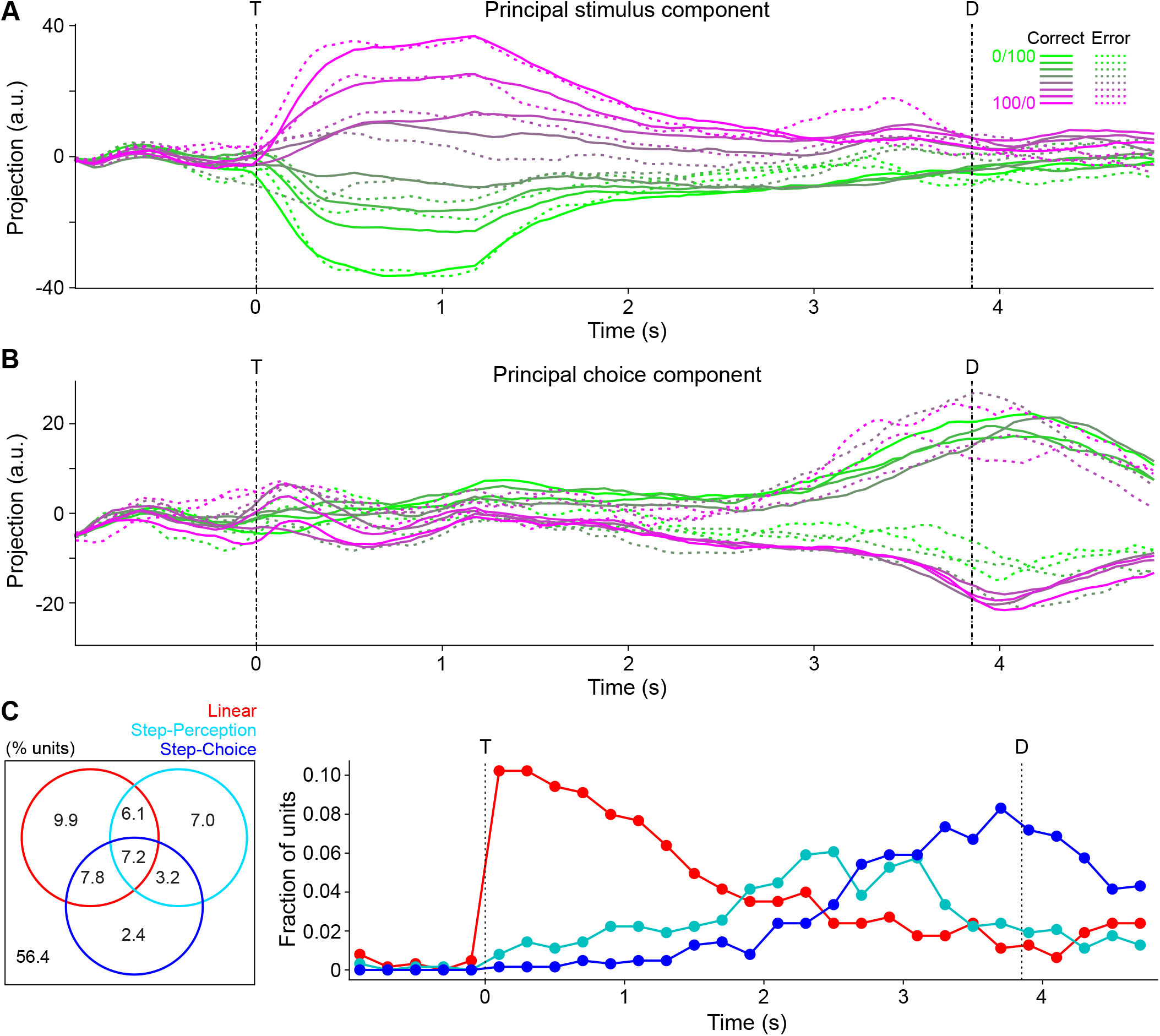
Modeled population activity and single unit coding properties: constrained units only. Compare with **Figures 6** and **S6**. A: Trial-averaged constrained pseudo-population activity projected onto demixed principal component of maximal stimulus-specific variance. Solid lines are correct trial averages; dotted lines are incorrect trial averages. Color scale corresponds to different mixture stimuli (%Sucrose/%NaCl). T: time of stimulus onset; D: decision time. **B**: Same as **A** but for the demixed principal component of maximal choice-specific variance. **C**: Left: Venn diagram showing percentages of constrained units (pooled over all models) with all possible combinations of coding types over time. Right: Distribution of coding types across constrained units over time.

**Supplementary Figure 6.**
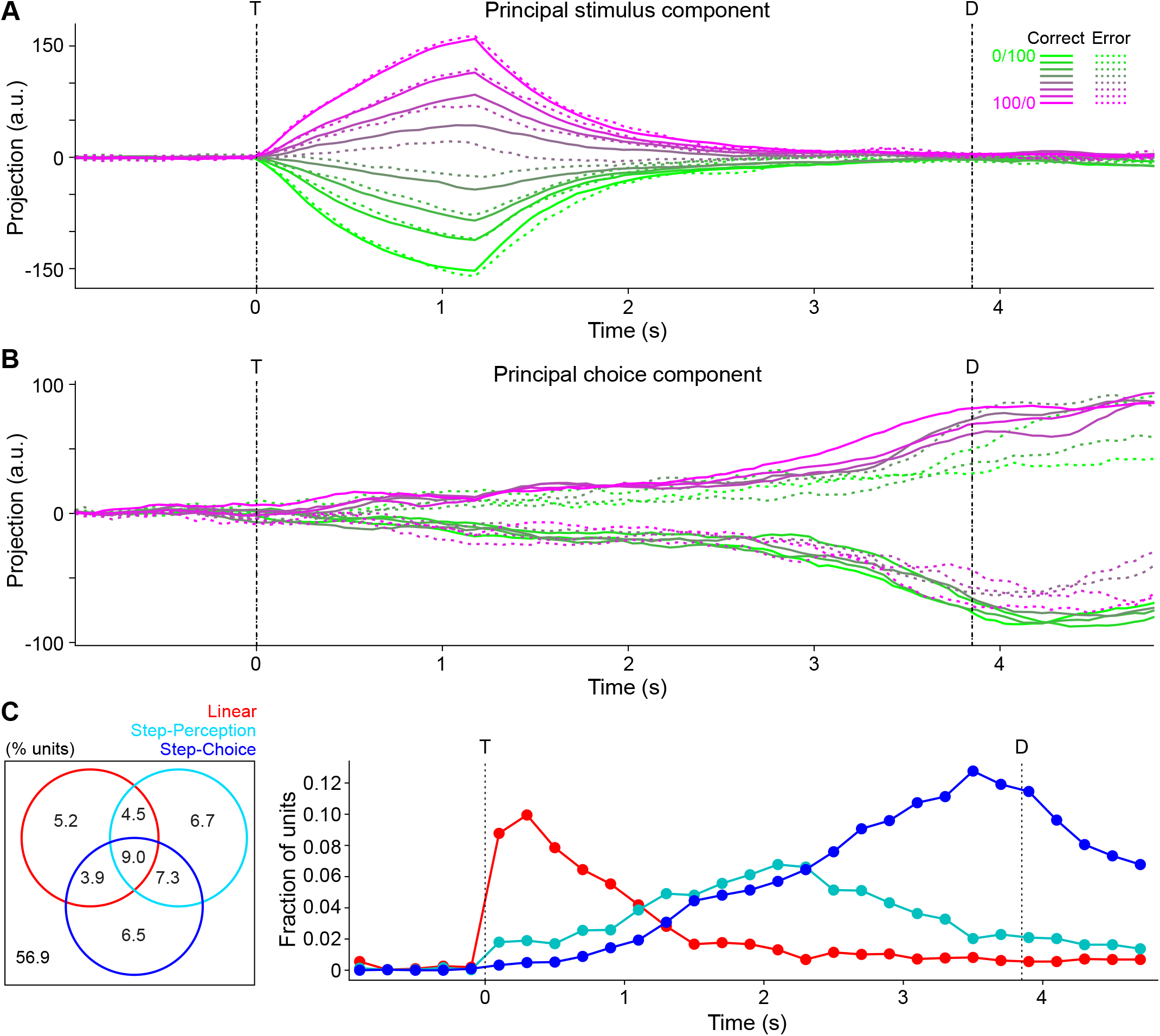
Modeled population activity and single unit coding properties: unconstrained units only. Compare with **Figures 6** and **S5. A**: Trial-averaged unconstrained pseudo-population activity projected onto demixed principal component of maximal stimulus-specific variance. Solid lines are correct trial averages; dotted lines are incorrect trial averages. Color scale corresponds to different mixture stimuli (%Sucrose/%NaCl). T: time of stimulus onset; D: decision time. **B**: Same as **A** but for the demixed principal component of maximal choice-specific variance. **C**: Left: Venn diagram showing percentages of unconstrained units (pooled over all models) with all possible combinations of coding types over time. Right: Distribution of coding types across unconstrained units over time.

**Supplementary Figure 7.**
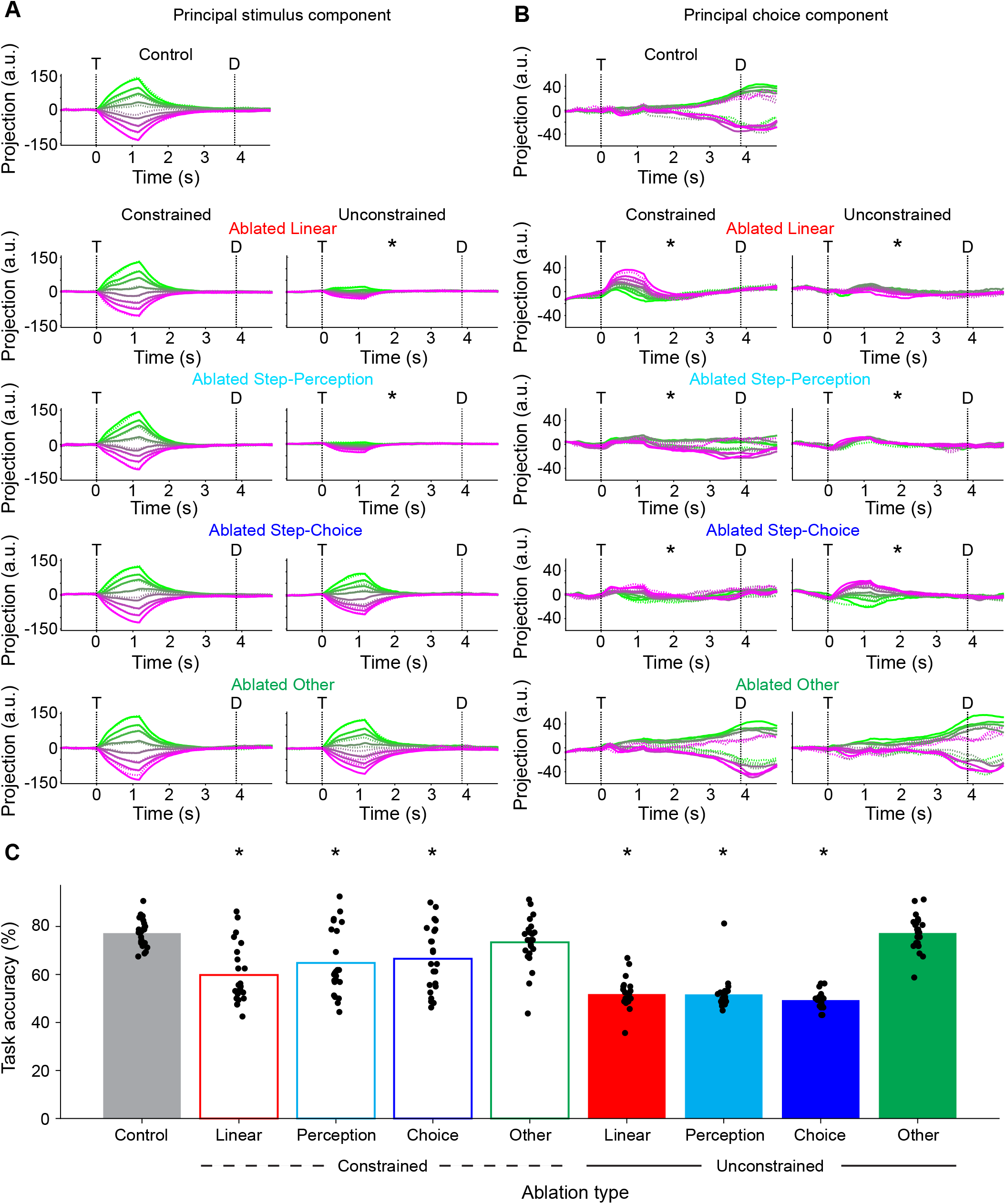
Effect of selective ablations on model dynamics and behavior: constrained vs unconstrained. **A-B**: Model dynamics after selectively ablating linear coding units, step-perception coding units, step-choice coding units, or “other” units in the constrained (left columns) or unconstrained (right columns) populations. Post-ablation pseudo-population activity is projected onto the stimulus (**A**) and choice-coding (**B**) components identified in the control condition (top). Color scale corresponds to different mixture stimuli (%Sucrose/%NaCl); solid and dashed lines correspond to correct and error trials. * indicates significant difference in mean absolute projections vs corresponding control condition (Dunnett’s test *p* < 0.01). T: time of stimulus onset; D: decision time. **C**: Behavioral performance of all models after selectively ablating categories of coding units. Bars represent means. * indicates significant difference in task accuracy vs control condition (Dunnett’s test *p* < 0.01).

**Supplementary Figure 8.**
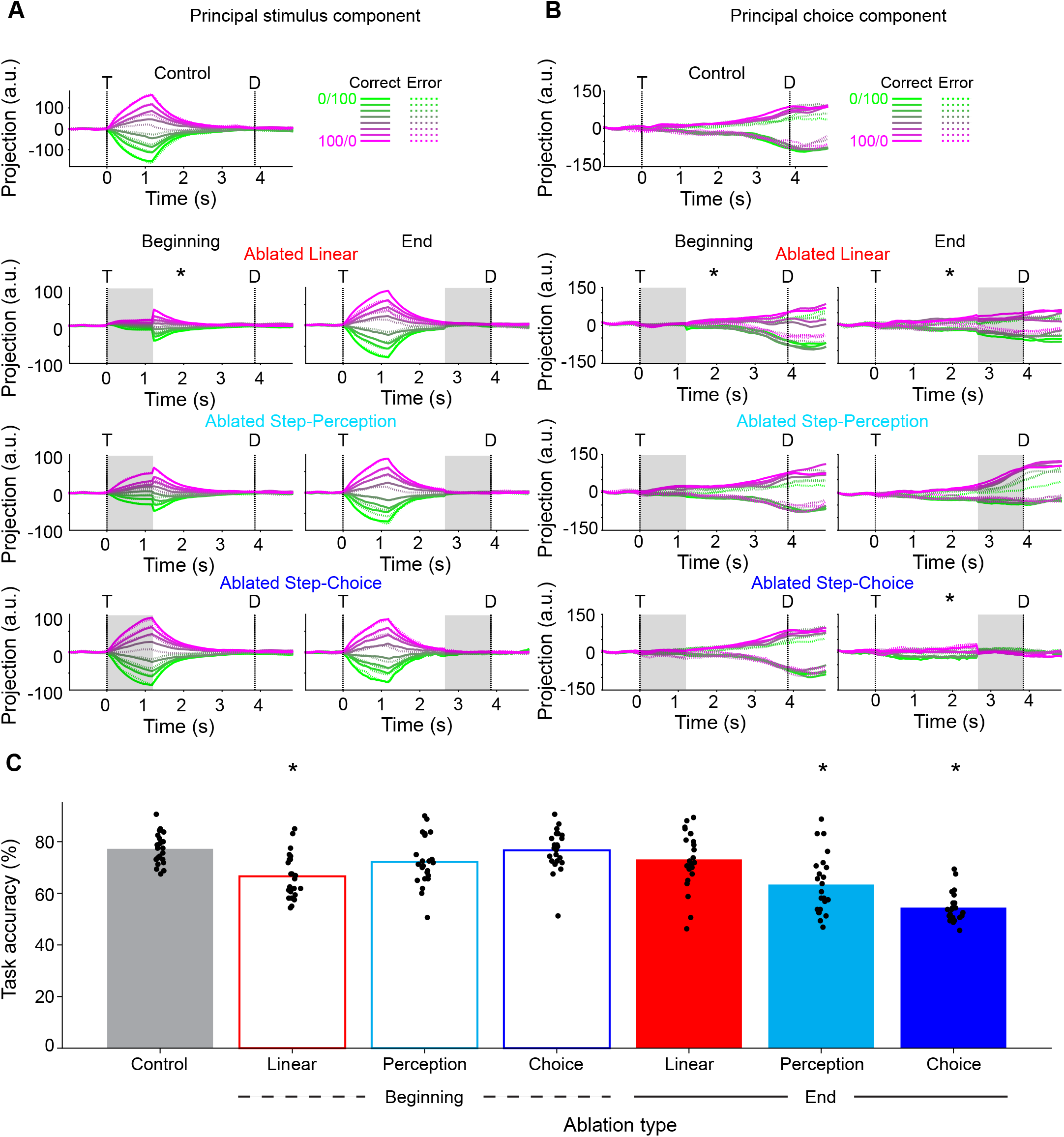
Effect of temporally restricted selective ablations on model dynamics and behavior: beginning vs end. **A-B**: Model dynamics after selectively ablating linear coding units, step-perception coding units, step-choice coding units, or “other” units at the beginning of the trial (left columns) or the end of the trial (right columns). Post-ablation pseudo-population activity is projected onto the stimulus (**A**) and choice-coding (**B**) components identified in the control condition (top). Color scale corresponds to different mixture stimuli (%Sucrose/%NaCl); solid and dashed lines correspond to correct and error trials. * indicates significant difference in mean absolute projections vs corresponding control condition (Dunnett’s test *p* < 0.01). T: time of stimulus onset; D: decision time. The beginning is [T, T + 1.2 s]; the end is [D – 1.2 s, D]. **C**: Behavioral performance of all models after selectively ablating categories of coding units in the beginning or end of the trial. Bars represent means. * indicates significant difference in task accuracy vs control condition (Dunnett’s test *p* < 0.01).

